# AMPK activation prevents hepatocellular carcinoma development through inhibition of HNF4α activity

**DOI:** 10.1101/2025.05.16.652569

**Authors:** Zhen Sun, Bernard Linares, Cassidy Urdiales, Fiyad Al-sarmi, Nagireddy Putluri, Jeanine L Van Nostrand

## Abstract

Hepatocellular carcinoma (HCC) is a major cause of cancer-related mortality and is largely driven by metabolic disorders such as obesity and Type 2 Diabetes. The AMP-activated protein kinase (AMPK) is a master regulator of metabolism, coordinating glucose and lipid metabolism, and its activation has been proposed as a therapeutic strategy for treating metabolic disorders. However, while AMPK activity has been reported to be downregulated in HCC, the precise role of AMPK in HCC development has not been clearly delineated. Here, we investigated the ability of AMPK activation to prevent HCC development using genetic models and specific allosteric AMPK activators. By leveraging a constitutively active AMPK transgenic mouse model and a pharmacological AMPK activator, we were able to elucidate the direct effects of AMPK activation on HCC development and progression. We observed that AMPK activation significantly reduced tumor formation in both diethylnitrosamine (DEN)-induced and streptozocin-induced (STAM) models of HCC. Our findings further implicate bile acid metabolism and hepatic nuclear factor alpha (HNF4α) signaling in the mechanism of AMPK-dependent HCC prevention. These findings provide mechanistic insights into AMPK biology and highlight the potential of AMPK as a therapeutic target, emphasizing the intricate interplay between metabolic dysregulation and cancer development.

## Introduction

Hepatocellular carcinoma (HCC) is a leading cause of cancer-related mortality worldwide, with its incidence rising sharply over the past few decades^1^. This increase is intricately linked to the growing prevalence of metabolic disorders such as obesity and Type 2 Diabetes, which significantly elevate the risk of HCC development^2^. The liver, being central to metabolic regulation, is particularly susceptible to cancer when metabolic homeostasis is disrupted^3^. Metabolically associated fatty liver disease (MAFLD), which can progress to metabolically associated steatohepatitis (MASH), fibrosis, cirrhosis, and ultimately HCC, exemplifies this connection^4^. The risk of developing HCC increases nearly 100-fold upon the onset of cirrhosis, underscoring the critical interplay between metabolic dysregulation and cancer susceptibility^5^.

Understanding how metabolic disorders increase the risk of HCC is important for addressing the rising rate of HCC-related deaths. The anti-diabetic medication metformin, a frontline treatment for Type 2 Diabetes, has shown promise in protecting against HCC^6–12^. Metformin use is associated with a lower incidence of HCC and extended overall survival in diabetic patients with HCC^9,10,13^. Furthermore, results from previous preclinical studies suggest that metformin influences the development of multiple cancers^14–16^. Mechanistically, metformin inhibits mitochondrial complex I, subsequently activating the energy stress sensor protein AMP-activated protein kinase (AMPK)^17–19^. However, some findings indicate that AMPK is not responsible for all of metformin’s beneficial effects, leaving the exact mechanism behind metformin’s tumor-suppressive properties unclear^19–23^.

AMPK is a crucial regulator of cellular energy homeostasis and is implicated in various metabolic processes^24–29^. AMPK operates downstream of the tumor suppressor liver kinase B1 (LKB1/*STK11*), the loss of which has been shown to lead to glucose intolerance and spontaneous tumor development in murine models^30–36^. Additionally, AMPK serves as a critical regulator of hepatic homeostasis, with low levels of phospho-AMPK in the liver correlating with an elevated risk of HCC development, aggressive pathological characteristics, and unfavorable prognoses in HCC patients^37–41^. These findings suggest that impaired AMPK activation may play a pivotal role in promoting HCC development and progression. Despite these correlative observations, definitive evidence demonstrating that direct AMPK activation can prevent HCC has been lacking. To address this, we directly investigated the role of AMPK in HCC prevention.

## Results

### AMPK activation prevents DEN model of HCC

To address the role of AMPK in HCC, we used the constitutively active AMPK mouse model, which harbors a truncated AMPKa1 protein (L-iAMPK^CA^; AMPK^CA^)^42^. In this model, expression of AMPK^CA^ is regulated by a lox-stop-lox and doxycycline-inducible promoter to provide both spatial and temporal control of expression (Figure S1A). These mice were bred to a mouse expressing Cre recombinase from an albumin promoter, such that recombination of the lox-stop-lox cassette, and corresponding AMPK activation, specifically occurs in hepatocytes of the liver^43^. Mice were fed a high fat diet (HFD) consisting of 45 kcal% fat and 17 kcal% sucrose containing doxycycline for the indicated amounts of time to induce AMPK^CA^ expression.

We first confirmed expression of the AMPK^CA^ transgene by performing western blot analysis of liver tissue from mice exposed ad libitum to HFD containing doxycycline for 7 days. A band at 32 kDa, corresponding to the truncated AMPK^CA^ transgene, was detected using both total AMPK and phospho-T172 AMPK antibodies. This band was observed exclusively in the livers of AMPK^CA^ mice, but not in wild-type (WT) mice carrying the transgene without albumin-Cre expression, despite receiving the same doxycycline-containing diet (Figure 1A). To assess AMPK activity, we examined phosphorylation of known AMPK substrates, including P-S792 RAPTOR and P-S79 ACC, by western blot under the same dietary conditions (Figure 1A). Additionally, we conducted immunohistochemical analysis for phospho-T172 AMPK and P-S79 ACC after 7 months of HFD with doxycycline to evaluate long-term pathway activation (Figure 1B)^44^. In both short-term and long-term settings, we observed robust activation of the AMPK signaling pathway in AMPK^CA^ livers compared to controls (Figure 1A–B).

**Figure 1.**
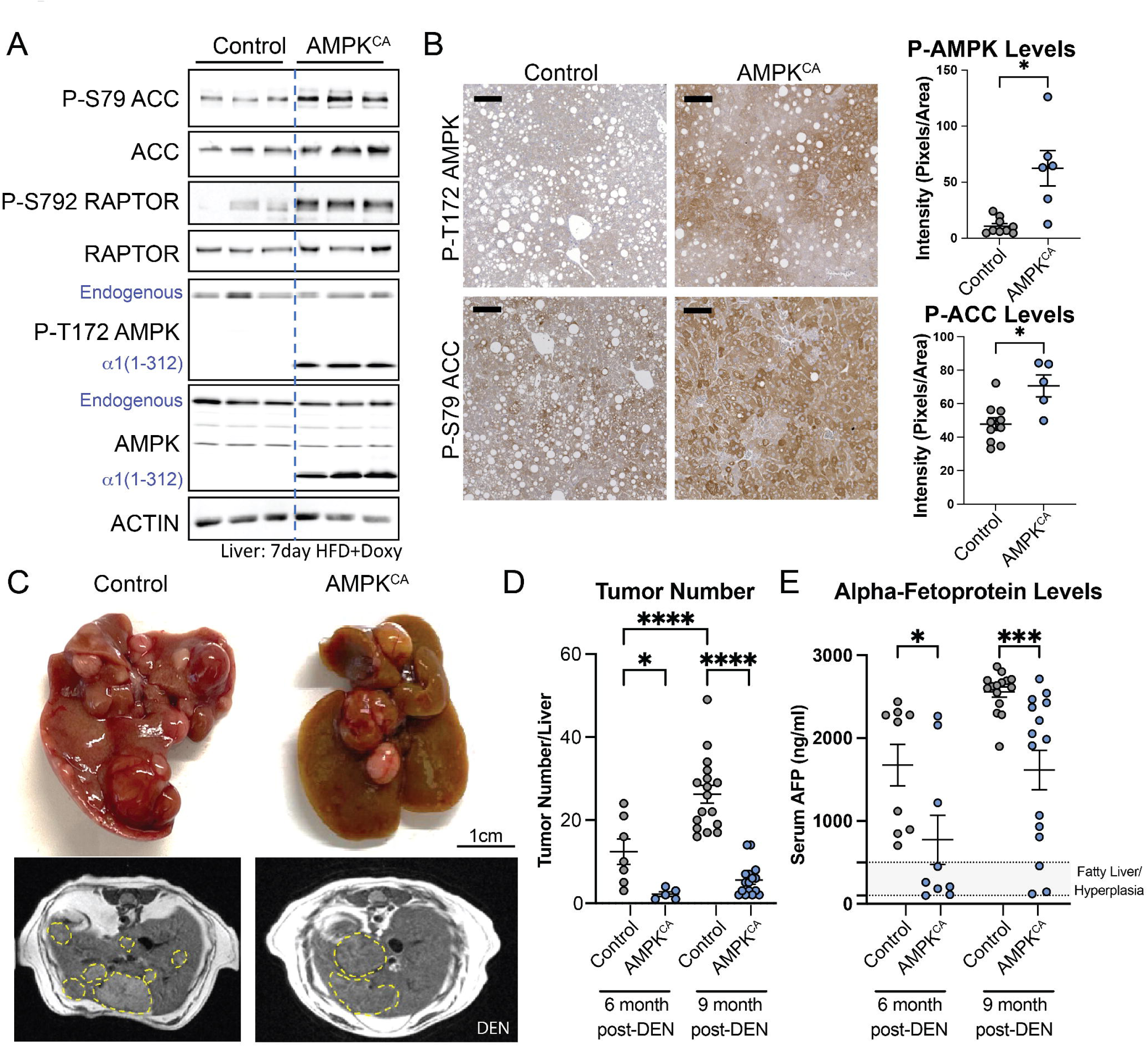
Genetic AMPK activation prevents HCC A. Western blot analysis of female mouse livers fed high fat diet (HFD) containing doxycycline (Doxy) to express constitutive active AMPK (a1(1-312); AMPK^CA^) for 7 days compared to control livers. B. Immunohistochemical analysis of livers from male control and AMPK^CA^ mice fed HFD+Doxy for 7 months. Scale bar 100µm. Right: Quantification of staining intensity for P-T172 AMPK and P-S79 ACC staining. n=6-10, ±SEM, Welch t-test. C. Representative whole-mount (top) and MRI (bottom) image of male control and AMPK^CA^ livers 9 months post-DEN injection. D. Quantification of tumor number in male control and AMPK^CA^ livers 6 months and 9 months post-DEN injection based on MRI images. 6m: N=5-7; 9m: N=16-17; Fisher LSD test. E. Quantification of serum alpha-fetoprotein (AFP) levels in control and AMPK^CA^ male mice 6 and 9 months post-DEN injection. n=9-15 Fisher LSD test. *<0.05, ***<0.001, ****<0.0001

To induce metabolic dysfunction, mice were fed HFD for an extended period. Consistent with previous findings, AMPK^CA^ mice exhibited increased resistance to diet-induced obesity (DIO) after 8 weeks of HFD with Doxycycline, as evidenced by a 12% reduction in body weight and improved glucose tolerance—an indicator of diabetes—compared to control mice (Figures S1B–D)^45^. Moreover, after 24 weeks on HFD, AMPK^CA^ mice displayed significantly lower hepatic triglyceride levels and lipid accumulation relative to control mice (Figure S1E-F), aligning with prior studies demonstrating that AMPK activation mitigates metabolic dysfunction and prevents the development of metabolic-associated fatty liver disease (MAFLD)^45^.

To investigate the impact of constitutively active AMPK in HCC development, we employed a diethylnitrosamine (DEN)-induced HCC model. Mice were administered DEN, a known liver carcinogen, at 2 weeks of age to initiate liver damage^46^. Beginning at 8-12 weeks of age, the mice were fed a HFD containing doxycycline to induce AMPK^CA^ expression under conditions of metabolic stress (Figure S1G). This timeline ensured that doxycycline administration did not interfere with DEN-induced mutagenesis or liver development. Mice were monitored for HCC progression using MRI from 6 to 9 months post DEN-injection, and livers were collected at 9 months – when control animals began to show signs of morbidity – for further analysis. Tumor number and size were quantified at both the 6- and 9-month timepoints, corresponding to the earliest detectable tumor burden and the study endpoint, respectively.

Between the 6- and 9-month timepoints, both control and AMPK^CA^ male mice exhibited a two-fold increase in tumor number, consistent with progressive tumor development. However, AMPK^CA^ mice had 83% and 79% fewer tumors than control littermates at 6 and 9 months, respectively, indicating a sustained suppression of tumor formation (Figures 1C–D, S1H). This effect was most pronounced in male mice, where HCC incidence is typically higher, but a similar trend was also observed in female mice at 9 months post-DEN injection (Figure S1I).

To validate the MRI findings, we measured serum alpha-fetoprotein (AFP) levels—a biomarker of HCC—at 6 and 9 months post-DEN treatment^47^. While AFP levels increased over time in both groups, levels in AMPK^CA^ mice remained significantly lower than in controls, suggesting a reduced tumor burden (Figure 1E). Collectively, these results support the conclusion that AMPK activation is sufficient to inhibit HCC development in the context of obesity.

### Tumors arise in the absence of AMPK activation

While AMPK activation was sufficient to suppress HCC tumor formation, we sought to understand the impact of AMPK activation in the tumors that did form. Surprisingly, despite the reduced tumor number in AMPK^CA^ mice (Figure 1D), the tumors that did form were, on average, significantly larger than those in control at the 9-month timepoint—a difference not observed at 6 months (Figure 2A). To determine whether the increased tumor size was due to enhanced proliferation, we assessed proliferative index. Mice were injected with BrdU 4 hours prior to tissue collection, and BrdU incorporation was assessed using immunohistochemistry. Minimal BrdU incorporation was detected in non-tumor liver tissue, consistent with the largely quiescent nature of the liver (Figure S2A). In contrast, tumors from AMPK^CA^ mice showed a marked increase in BrdU incorporation and a higher number of mitotic figures compared to control tumors (Figure 2B, S2B). These findings suggest that the rapid increase in tumor size between 6 and 9 months in AMPK^CA^ mice is at least partially driven by increased proliferative capacity.

**Figure 2.**
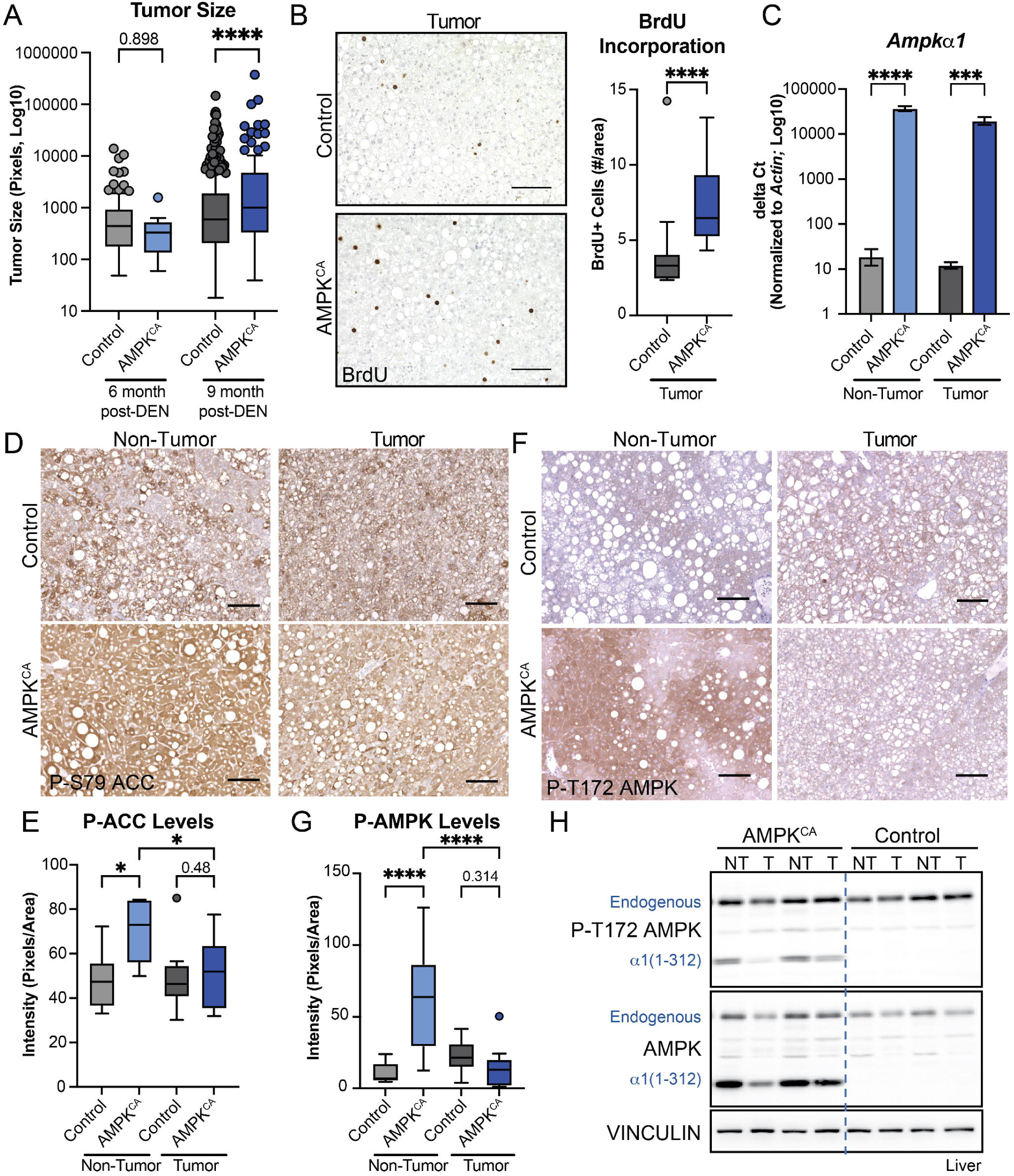
Tumors arise in the absence of AMPK Activity A. Quantification of tumor size from MRI imaging at 6 and 9 months. n=10-448 Fisher LSD test. B. Immunohistochemical analysis of BrdU incorporation in tumor tissue from control and AMPK^CA^ livers following 4-hour BrdU pulse (10mg/kg). Scale bar 100µm. Right: Quantification of number of positive cells per area. n=13-17. Welch t-test. C. Gene Expression of Ampka1 normalized to *Actin* in non-tumor and tumor tissue. n=5-6. ±SEM, Fisher LSD test D. Immunohistochemical analysis of P-S79 ACC in non-tumor and tumor tissue from control and AMPK^CA^ livers. Scale bar 100µm. E. Quantification of staining intensity of P-S79 ACC. n=5-14. Fisher LSD test. F. Immunohistochemical analysis of P-T172 AMPK in non-tumor and tumor tissue from control and AMPK^CA^ livers. Scale bar 100µm. G. Quantification of staining intensity. n=6-11. Fisher LSD test. H. Western blot analysis of non-tumor (NT) and tumor (T) tissue. VINCULIN as loading control. *<0.05, **<0.01, ***<0.001, ****<0.0001

This raised the possibility that tumors either escaped AMPK activation or that AMPK activation could paradoxically promote tumor growth once tumors had formed. To assess the role of AMPK activation within the tumors, we first evaluated whether the tumors originated from cells that had escaped Cre-mediated recombination of the lox-stop-lox cassette by examining the expression of mKate, a fluorescent marker downstream of the recombined lox site, in dissected tumors and adjacent non-tumor tissue (Figures S1A,S2C). High levels of mKate expression were observed in both non-tumor and tumor tissues, indicating successful recombination. We next assessed the tumors’ responsiveness to doxycycline by evaluating expression of Gfp, which is driven by the tet-responsive promoter. Similar to mKate, Gfp expression was robust in both non-tumor and tumor tissues of AMPK^CA^ mice but absent in wild-type controls (Figures S2D). Finally, we evaluated AMPK^CA^ transgene expression by measuring AMPKα1 mRNA levels. Tumors exhibited high AMPKα1 transgene expression, comparable to adjacent non-tumor tissues (Figure 2C). Together, these results indicate that the tumors did not escape Cre recombination or doxycycline responsiveness and maintained AMPK transgene expression at the mRNA level.

We next assessed AMPK activity by western blot and immunohistochemical analysis using P-S79 ACC and P-T172 AMPK as markers. Although both markers were elevated in non-tumor liver tissue from AMPK^CA^ mice relative to controls, AMPK activity was markedly reduced within the tumors compared to matched non-tumor tissue (Figures 2D–H, S2E). In addition, total AMPK^CA^ protein levels were decreased in 5 out of 8 tumors relative to adjacent non-tumor tissue (Figure 2H). Notably, in 3 of these tumors, endogenous AMPK protein levels were also reduced, suggesting a broader suppression of AMPK protein expression. This reduction in AMPK protein and activity was despite comparable levels of transgenic GFP protein in non-tumor and tumor tissue (Figure S2F). These observations indicate that tumors actively downregulate AMPK activity and protein levels to enable tumor development and growth, highlighting a mechanism by which tumors may evade the tumor-suppressive effects of AMPK activation, ultimately contributing to HCC development and progression.

### AMPK activation prevents STAM model of HCC

To further investigate AMPK’s role in preventing HCC, we employed the STAM model, which recapitulates features of diabetes-associated HCC. In this model, mice were injected with the pancreatic toxin streptozotocin (STZ) at postnatal day 2 to induce Type I Diabetes^48^. Beginning at 4 weeks of age, mice were fed HFD containing doxycycline to promote fatty liver, fibrosis, and ultimately HCC by 16 weeks of age, with livers collected at 20 weeks for tumor analysis (Figure S3A)^48^. At 8 weeks, body weights were comparable between control and AMPK^CA^ mice; however, by 16 weeks, AMPK^CA^ mice exhibited a modest but statistically significant reduction in body weight (Figure 3A).

**Figure 3.**
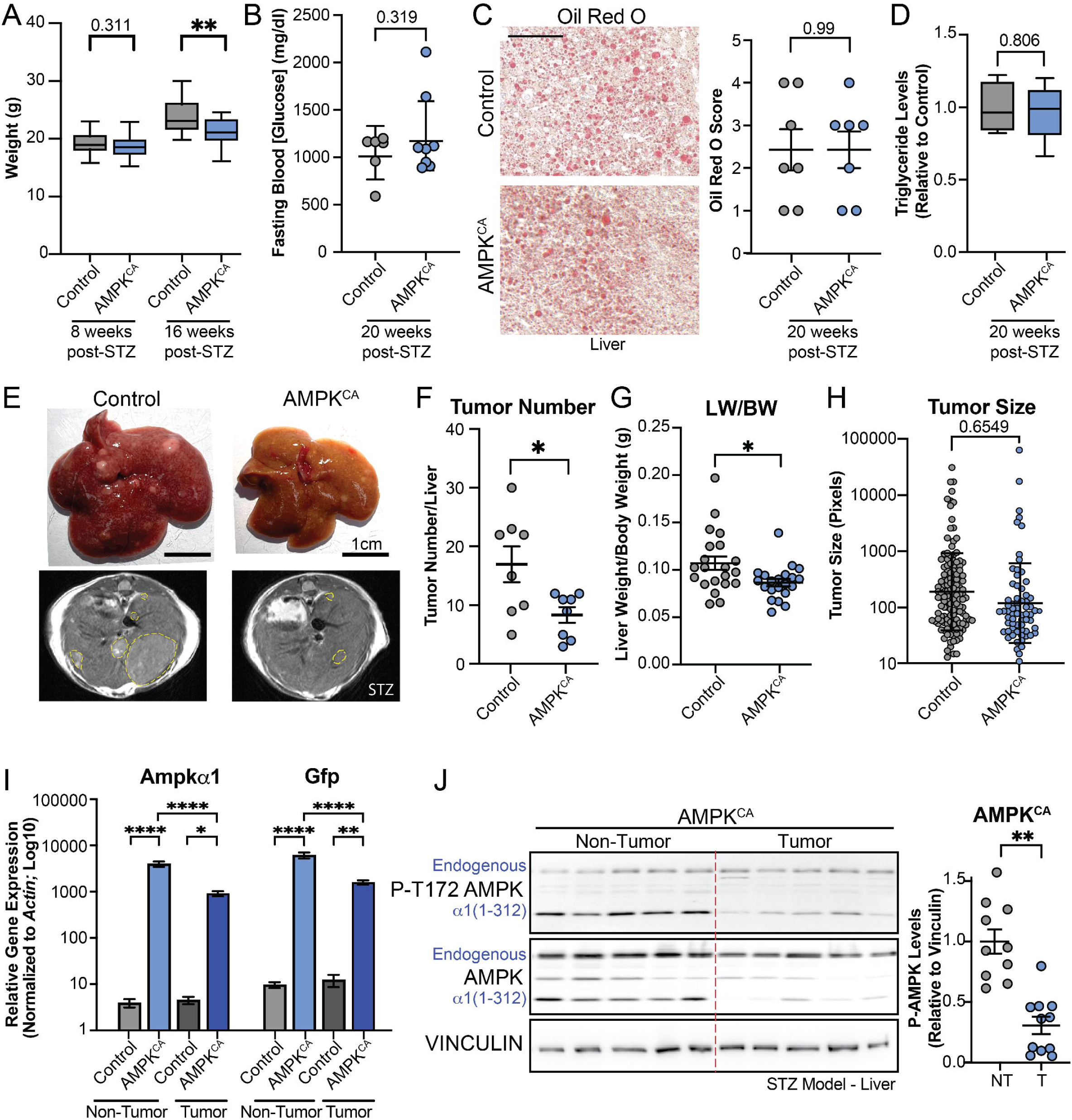
Genetic AMPK Activation prevents HCC in STAM model A. Weight of male control and AMPK^CA^ mice on high fat diet (HFD) for 8 and 16 weeks following STZ injection n=14-19. Fisher LSD test B. Fasting blood glucose levels 20 weeks post-STZ injection. n=6-8. Welch t-test, C. Oil Red O staining of control and AMPK^CA^ livers 20 weeks following STZ injection. Right: Scoring of Oil Red O levels, n=7. Welch t-test Scale bar 100µm. D. Quantification of triglyceride levels in livers of control and AMPK^CA^ mice 20 weeks post-STZ injection. n=5. Welch t-test. E. Whole-mount image (top) and MRI image (bottom) of livers 20 weeks following STZ injection. F. Quantification of tumor number from control and AMPK^CA^ mice injected with STZ. n=8. Welch t-test. G. Liver weight relative to body weight of STZ-injected mice n=13-14. Welch t-test. H. Quantification of tumor sizes 20 weeks following STZ injection n=67-136. Welch t-test I. Gene expression analysis of non-tumor and tumor tissue from STZ-injected mice n=6-7. Fisher LSD test. J. Western blot analysis of matched non-tumor and tumor tissue from AMPK^CA^ mice injected with STZ, Right: Quantification of P-AMPK transgene levels, n=10-11 Welsh t-test. *<0.05, **<0.01, ***<0.001, ****<0.0001

To assess metabolic function, we measured fasting blood glucose levels at 8 and 20 weeks post-STZ injection. Both genotypes showed elevated glucose levels, consistent with impaired pancreatic function (Figures 3B, S3B). No significant differences were observed between control and AMPK^CA^ mice, suggesting that hepatic AMPK activation alone is insufficient to restore systemic glucose homeostasis in this model. Given the presence of metabolic-associated steatotic liver disease (MASLD) in this model, we next assessed hepatic lipid accumulation. Oil Red O staining of liver sections at 20 weeks post-STZ injection revealed no discernible differences in lipid content between control and AMPK^CA^ mice (Figure 3C). This finding was supported by lipidomics analysis, which showed comparable hepatic triglyceride levels across genotypes (Figure 3D), indicating that AMPK activation in the liver does not reduce lipid accumulation in the STZ model.

We then examined the impact of AMPK activation on HCC development. Using MRI, we quantified tumor number and size. Consistent with findings from the DEN model, AMPK^CA^ mice exhibited a significant reduction in tumor burden relative to controls (Figures 3E–F). This reduction was also reflected in decreased liver weight-to-body weight ratios in AMPK^CA^ mice (Figure 3G).

Additionally, we observed a trend toward smaller tumor sizes in AMPK^CA^ mice, resembling our observations in the earlier 6-month timepoint in the DEN-induced model (Figure 3H).

To evaluate transgene expression and AMPK activity in tumors, we performed gene expression analysis of Ampkα1 and Gfp, as well as western blot analysis of macroscopically dissected tumor and non-tumor tissue (Figures 3I–J). Similar to the DEN model, tumors from AMPK^CA^ mice expressed both Ampkα1 and Gfp at levels comparable to adjacent non-tumor tissue (Figure 3I). However, AMPK activity and protein levels were markedly reduced in tumor tissues (10 out of 11 tumors), suggesting suppression of AMPK signaling (Figures 3I–J). Consistent with our findings in the DEN model, endogenous AMPK protein levels were also diminished in 4 out of 5 tumors. These results collectively demonstrate that genetic activation of AMPK suppresses HCC development in the STAM model. Importantly, this tumor-suppressive effect occurs independently of improvements in glucose or lipid homeostasis, further emphasizing AMPK’s direct role in inhibiting tumorigenesis.

### Bile acid levels are increased upon AMPK activation

To understand how AMPK suppresses HCC formation, we performed targeted profiling of approximately 350 metabolites in livers from control and AMPK^CA^ mice^49^. We initially focused on evaluating metabolites in liver tissue from mice fed HFD or chow diet containing doxycycline at 2 timepoints of AMPK expression (15 days, 2 months). We extended these analyses to understand what metabolites were changed upon tumor formation by comparing metabolites in tumor and adjacent non-tumor tissue from matched livers 9 months post-DEN injection in the presence of HFD containing doxycycline for 7 months (Figure S4A). This comparison allows us to discern which metabolites were lost or gained upon tumor formation, suggesting a role for them in tumor prevention or initiation, respectively. From these analyses, we identified approximately 200 metabolites, of which 66 changed significantly in at least one condition based on average fold change and p-value calculations derived from 4-6 samples per genotype per timepoint (Figure S4A). Interrogation of metabolites that significantly changed in AMPK^CA^ tissue relative to control liver tissue identified increases in nucleotide and methionine metabolites, and decreases in glycolysis and TCA cycle intermediates, supporting AMPK’s role in metabolic rewiring (Figures S4B–C)^27^. These findings are consistent with previous results that AMPK activation suppresses catabolic processes, such as DNA synthesis, and increases anabolic processes, such as oxidative phosphorylation^50,51^.

We were interested in understanding what metabolites were consistently dependent on AMPK and maintained throughout tumor development. Therefore, we identified which metabolites were significantly altered in at least three of the four non-tumor conditions. From this analysis, only 5 metabolites, including malic acid, betaine aldehyde, taurolithocholic acid (TLCA), taurodeoxycholic acid (TDCA), and Glycohyodeoxycholic acid (GHDCA), were found to be enriched across multiple conditions and timepoints (Figure S4D). The identification of multiple bile acids (TLCA, TDCA, GHDCA) made bile acid metabolism an intriguing candidate for further investigation^52,53^. Quantification of total bile acids in the livers of AMPK^CA^ mice at 2 months following high fat diet and doxycycline administration revealed a significant increase in bile acid levels relative to control livers (Figures 4A, S4E). A more detailed evaluation of individual bile acids across timepoints revealed a marked increase in bile acids at most timepoints. Notably, the levels of total bile acids were decreased in the tumors relative to matched non-tumor tissue following 7 months of doxycycline treatment (corresponding to 9 months post-DEN injection), suggesting their levels were downregulated upon formation of the tumors, supporting their potential role in tumor prevention (Figure 4B).

**Figure 4.**
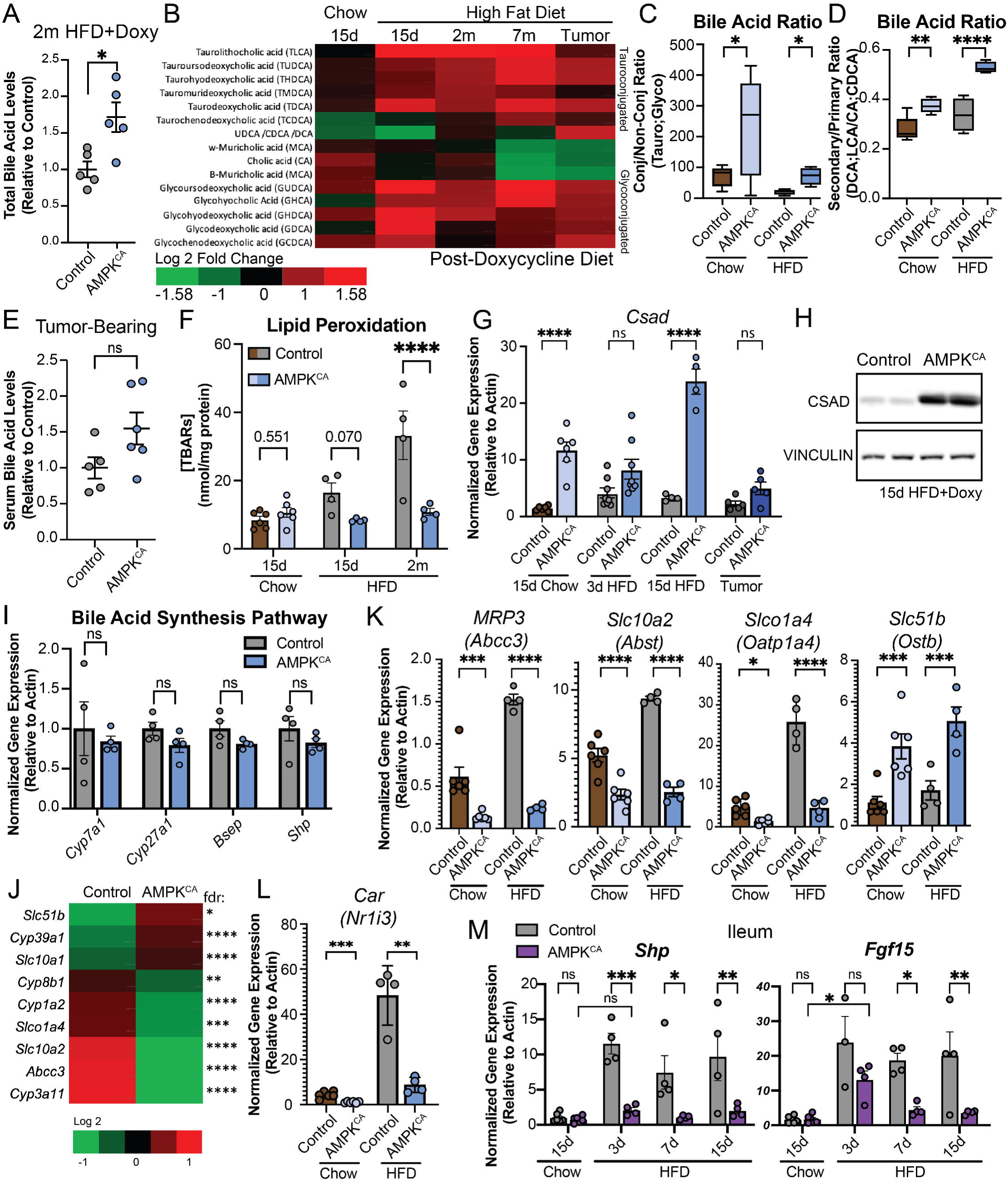
Bile Acid levels are increased due to transporter misregulation A. Quantification of relative levels of bile acids in livers from control and AMPK^CA^ mice following 2 months of doxycycline. n=5. Welsh t-test. B. Heatmap of average relative bile acids levels of AMPK^CA^ livers relative to control livers following varying durations of doxycycline treatment (15d, 2 month (2m)) or in matched non-tumor and tumor tissues (7 month (7m) vs Tumor), n=4-6/genotype per timepoint. C. Ratio of conjugated vs non-conjugated bile acids in livers from control or AMPK^CA^ mice on chow or high fat diet (HFD) for 15 days. n=4-6. Welsh t-test. D. Ratio of secondary (DCA;LCA) vs primary (CA;CDCA) bile acids in livers from control or AMPK^CA^ mice on chow or HFD for 15 days. n=4-6. Fisher LSD test. DCA: Deoxycholic acid, LCA: Lithocholic acid, CA: Cholic Acid, CDCA: Chenodeoxycholic acid. E. Quantification of serum bile acid levels relative to control livers of tumor bearing mice on doxycycline diet for 7 months. n=5-6. Welch t-test. F. Quantification of TBARS concentration in livers from control or AMPK^CA^ mice on chow or HFD containing doxycycline for 15 days or 2 months. n=4-6. Fisher LSD test. G. Quantification of gene expression in control or AMPK^CA^ livers of mice on chow or HFD containing doxycycline for 3 days or 15 days, or in tumors. n=4-6. Fisher LSD test. H. Western blot analysis of livers from mice on HFD containing doxycycline for 15 days. I. Gene expression of bile acid biosynthesis proteins and bile acid response genes in livers from mice fed HFD containing doxycycline for 15 days. n=4. Fisher LSD test. J. Heatmap of log2 fold change of gene expression changes in AMPK^CA^ relative to control livers from mice on HFD containing doxycycline for 2 months. K. Gene expression of bile acid transporters in livers from mice fed chow or HFD containing doxycycline for 15 days. n=4-6. Fisher LSD test. L. Gene expression of bile acid transcription factor in livers from mice fed chow or HFD containing doxycycline for 15 days. n=4-6. Fisher LSD test. M. Gene expression of bile acid responsive genes in ileum from mice fed HFD containing doxycycline for 15 days. n=4 Fisher LSD test. ±SEM *<0.05, **<0.01, ***<0.001, ****<0.0001

Bile acids are synthesized in the liver from cholesterol and play a key role in emulsifying dietary fats for absorption, while also acting as signaling molecules that regulate metabolism, inflammation, and liver regeneration^53^. They are secreted into the bile, modified by gut microbes into secondary bile acids, and recirculated back to the liver through the enterohepatic circulation^54^. In the liver, bile acids can be further conjugated with glycine or taurine, a process that increases their solubility and reduces cytotoxicity, enabling efficient secretion into bile and effective emulsification of dietary fats in the intestine^55^. Thus, we interrogated the relative status of the bile acids in the livers of AMPK^CA^ and control mice. Conjugated bile acids were the predominant form of bile acids increased, with the ratio of conjugated to non-conjugated significantly upregulated (Figure 4C). Taurine-conjugated bile acids accounted for much of these changes, with taurolithocholic acid (TLCA) being the most consistent and significantly upregulated bile acid across timepoints but being suppressed in the tumor. Of note, glycine-conjugated bile acids were also significantly upregulated, but overall levels were lower because glycine-conjugated bile acids are not the predominant form in mice (Figure 4B). We also observed an increase in the ratio of secondary to primary bile acids in AMPK^CA^ mice regardless of conjugation status (Figure 4D). Of note, our bile acid detection method was unable to distinguish between non-conjugated ursodeoxycholic acid, chenodeoxycholic acid, and deoxycholic acid preventing us from using these isoforms for comparing between primary and secondary bile acids^56^. However, the majority of changes in bile acids were driven by taurine-conjugated forms precluding the need for this comparison.

Previous publications have suggested that increased bile acid levels in the serum are correlated with a higher incidence of HCC in patients, suggesting that higher serum bile acid levels either support HCC development or serve as a biomarker of HCC^57^. Therefore, we compared the bile acid levels of control and AMPK^CA^ mice bearing HCC after 7 months of doxycycline (9 months post-DEN treatment). Intriguingly, despite observing an increase in bile acids in the non-tumor tissue at this timepoint, we did not observe a significant difference in serum bile acid levels between control and AMPK^CA^ mice (Figure 4E). This suggests that in our system bile acids in the serum do not directly correlate with bile acid levels in the liver.

In the liver, bile acid species have been shown to have contrasting roles in liver pathogenesis, largely involving effects on oxidative stress^58–62^. For example, bile acids have been attributed with both antioxidant properties in the liver (i.e., ursodeoxycholic acid – UDCA) as well as with the increase in lipid peroxidation and oxidative stress resulting in hepatotoxicity (i.e., chenodeoxycholic acid – CDCA)^56,63,64^. Therefore, we interrogated the levels of lipid peroxidation by evaluating the levels of thiobarbituric acid reactive substances (TBARS) in the liver upon the various timepoints of AMPK activation (Figure 4F)^58^. While we observed a marked increase in lipid peroxidation levels in the control livers following administration of HFD containing doxycycline, this increase was not observed in AMPK^CA^ livers (Figure 4F). We also observed no change in glutathione usage as determined by the GSH/GSSG ratio in livers at 2 months following AMPK activation in the presence of high fat diet (Figure S4F). Evaluation of previously published transcriptomic data of AMPK^CA^ mice following 2 months of HFD also indicated a decrease in glutathione transferase gene expression relative to control mice (Figure S4G)^45^. Thus, the increase in bile acids present in the liver is not resulting in an oxidative environment and AMPK activation is protective against HFD-induced oxidative stress.

The pro-oxidative and cytotoxic properties of bile acids are strongly influenced by their hydrophobicity; conjugation with amino acids such as taurine reduces this hydrophobicity, resulting in more water-soluble and less toxic bile salts^65–67^. In line with this, we observed increased expression of cysteine sulfinic acid decarboxylase (CSAD)—the rate-limiting enzyme in taurine biosynthesis—at both the mRNA and protein levels following AMPK activation in both the DEN and STZ models (Figures 4G,H,S4H)^68^, suggesting increased production of taurine. This upregulation may underlie the striking increase in taurine-conjugated bile acids, particularly TLCA, in AMPK^CA^ livers and points a potential mechanism by which AMPK can promote a less oxidative hepatic environment in the context of high fat diet. Notably, *Csad* expression was suppressed in tumor tissue from AMPK^CA^ mice, and low CSAD levels are associated with worse outcome of liver cancer patients (Figure S4I), supporting a role of taurine-dependent bile acid conjugation in the prevention of HCC.

To determine how AMPK activation leads to elevated hepatic bile acid levels, we analyzed the expression of key enzymes in the classical bile acid biosynthesis pathway important for the conversion of cholesterol into primary bile acids—*Cyp7a1*, *Cyp27a1*—two weeks following doxycycline administration under HFD conditions^69^. Unexpectedly, despite the elevated bile acid levels in AMPK^CA^ livers, there were no changes in the expression of these enzymes compared to control (Figure 4I). We also examined the activity of the farnesoid X receptor (FXR), a nuclear receptor activated by bile acids to maintain homeostasis, by measuring expression of its target genes *Shp* (*Nr0b2*) and *Bsep* (*Abcb11*)^70^. These genes are typically upregulated in response to high bile acid levels to prevent further accumulation and are reported to be negatively regulated by AMPK^71^. However, we observed no significant difference in expression of these genes, suggesting that FXR signaling is not activated in response to increased bile acid levels in AMPK^CA^ livers (Figure 4I).

To investigate how bile acid levels are altered in AMPK^CA^ livers, we analyzed the previously published RNA-seq data to assess changes in genes involved in bile acid metabolism and transport following AMPK activation^42^. This analysis revealed differential expression of several genes related to bile acid synthesis and transport (Figure 4J). Specifically, we observed upregulation of the alternative bile acid synthesis gene *Cyp39a1*, alongside downregulation of classical pathway genes *Cyp8b1* and *Cyp3a11* (the mouse ortholog of human *CYP3A4*)^53,72–76^. Elevated *Cyp39a1* expression has been associated with reduced HCC risk, while *Cyp8b1* deficiency prevents diet-induced obesity and improves, suggesting potential protective roles in AMPK^CA^ livers^77–80^. Meanwhile, *Cyp3a11* is typically upregulated in response to bile acid toxicity, and its downregulation here may indicate that AMPK^CA^ livers are not activating the liver’s adaptive detoxification mechanisms^81–83^. However, the expression changes observed in these bile acid synthesis genes alone, which do not involve rate-limiting steps, are unlikely to account for the overall elevated bile acid levels seen in AMPK^CA^ livers.

In contrast, changes in bile acid transporters were more striking. We observed downregulation of several bile acid exporters, including *Abcc3* (*Mrp3*), *Slc10a2* (*Asbt*), and *Slco1a4* (*Oatp1a4*, the homolog of human *OATP1A2*), along with upregulation of import-associated genes such as *Slc10a1 (Ntcp)* and *Slc51b (Ostb)*^64,75^. Among these, *Abcc3* stood out as one of the most significantly downregulated genes. Quantitative RNA analysis confirmed marked suppression of *Abcc3* transcript levels in both chow-fed and HFD-fed AMPK^CA^ mice (Figure 4K). This downregulation was already evident three days after initiating doxycycline treatment (Figure S4J), indicating that it may be a direct consequence of AMPK activation. We further evaluated expression of the other bile acid and bile salt transporters. Consistent with the RNA-seq, we observed decreased expression of *Slc10a2* and *Slco1a4*, both of which mediate bile acid transport from hepatocytes to cholangiocytes. In contrast, *Slc51b* (*OSTβ*), which can facilitate both bile acid import and export, was upregulated providing a potential avenue for increasing bile acid levels (Figure 4K). Similar transporter expression patterns were also observed in chow-fed

AMPK^CA^ mice and in the STAM model, where AMPK^CA^ mice develop comparable levels of fatty liver, suggesting these changes are not simply a response to hepatic lipid accumulation (Figure S4K). Notably, we also observed reduced expression of the nuclear receptor *CAR* (*Nr1i3*, constitutive androstane receptor), a key regulator of genes involved in bilirubin clearance, including *Cyp8b1* (Figure 4L)^84^. Collectively, these data suggest that altered expression of bile acid transporters—particularly reduced efflux capacity—likely contributes to the accumulation of bile acids in AMPK^CA^ livers. Furthermore, the observed increase in the ratio of secondary to primary bile acids supports a model in which bile acids are efficiently imported into the liver but not effectively exported.

To determine whether the elevated hepatic bile acid levels observed in AMPK^CA^ mice, despite the reduced expression of bile acid exporters, led to increased systemic bile acid levels via the enterohepatic circulation, we examined the expression of bile acid-responsive genes *Shp* and *Fgf15* in the distal ileum^71^. In control mice, administration of a HFD triggered a rapid increase in *Shp* and *Fgf15* expression within three days compared to chow-fed mice. In contrast, AMPK^CA^ mice showed only a modest increase in *Shp* and *Fgf15* expression at day 3, which was diminished by day 7 (Figure 4M). These findings suggest that the accumulation of hepatic bile acids in AMPK^CA^ mice does not induce a systemic response through the intestinal arm of the enterohepatic circulation. This supports the notion that enterohepatic signaling is disrupted, likely due to impaired bile acid export from the liver. Collectively, these results indicate that AMPK activation alters hepatic bile acid composition and retention in a manner that may be protective, even under high-fat diet conditions.

### AMPK inhibits HNF4α activity

To understand how AMPK influences bile acid metabolism, we analyzed transcriptional changes in liver tissue from AMPK^CA^ mice using published RNA-seq data^42^. This analysis identified 286 upregulated and 361 downregulated genes compared to controls. Pathway enrichment analysis showed that upregulated genes were significantly associated with xenobiotic metabolism, bile acid metabolism, oxidative phosphorylation, and fatty acid metabolism. In contrast, downregulated genes were enriched in pathways related to epithelial-to-mesenchymal transition, interferon responses, and cholesterol homeostasis (Figure S5A). Notably, many of the most significantly altered genes were directly involved in bile acid regulation (Figure S5B). The enrichment of bile acid and xenobiotic metabolism pathways—both of which are involved in the modification and clearance of bile acids—supports the conclusion that AMPK activation significantly alters bile acid handling in the liver.

Next, we explored how AMPK activation modulates the expression of bile acid transporters by analyzing transcription factors associated with the observed gene expression changes. Pathway analysis identified transcription factors linked to oxidative stress responses, lipid metabolism, and glucose homeostasis—such as NRF2, USF1/2, and PPARD—as well as inflammatory regulators like RELA, NFKB1, and STAT3 (Figures 5A, S5C). Many of these factors have previously been associated with gene expression changes following AMPK activation^18,85–87^. Among the most significantly enriched regulators was HNF4α, a key transcription factor governing hepatocyte identity and bile acid metabolism^88–90^ (Figure 5A, S5C). Gene set enrichment analysis further revealed a negative correlation between HNF4α target genes and those upregulated in AMPK^CA^ liver (Figure 5B). Based on these findings, we focused subsequent analysis on the role of HNF4α in AMPK-mediated regulation of bile acids and HCC prevention.

**Figure 5.**
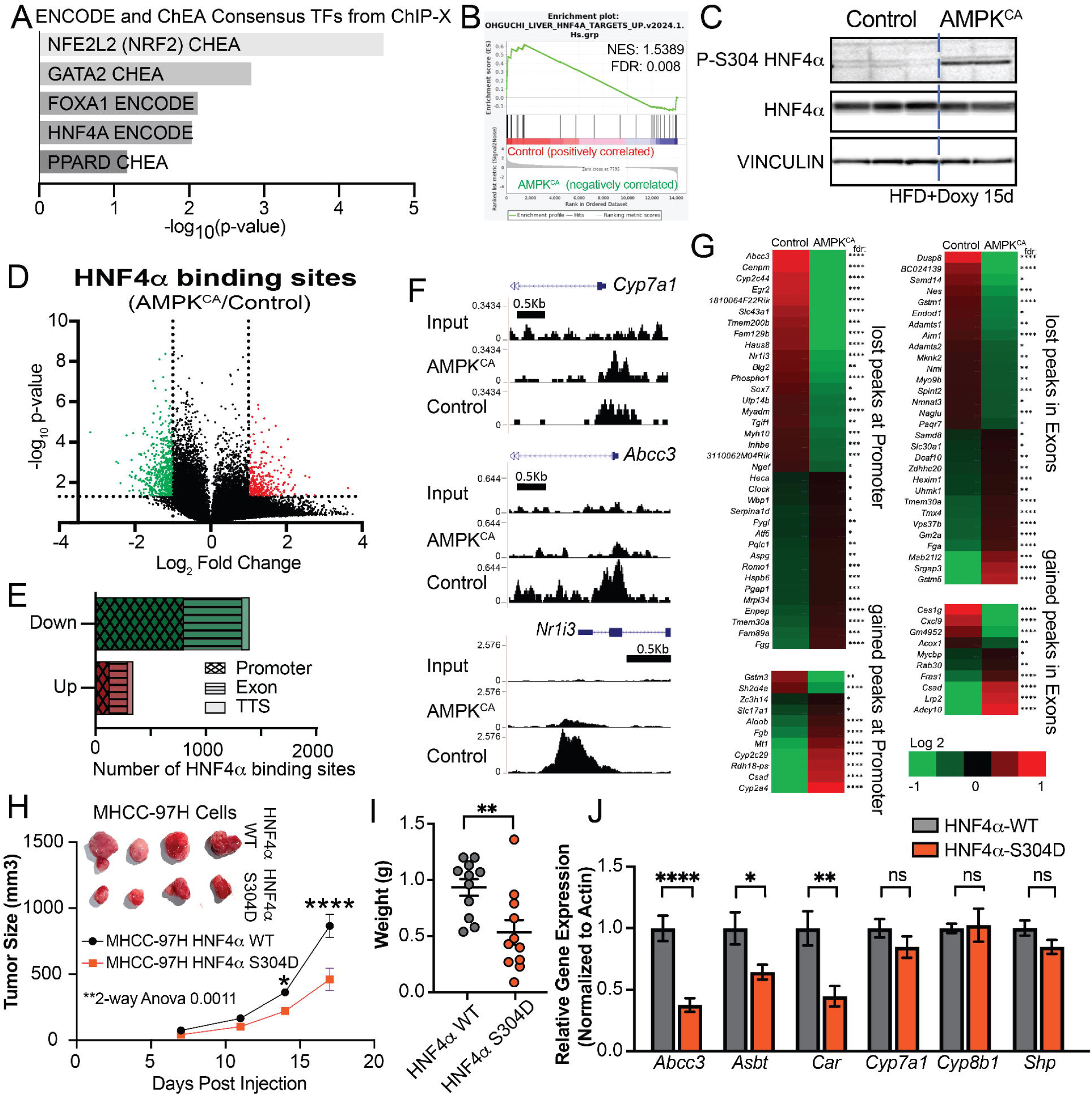
AMPK regulates HNF4α to drive tumor suppression A. Transcription factors enriched in AMPK^CA^ mice relative to control mice following 2 months of HFD containing doxycycline using Enrichr and a 1.3 fold change with an adjusted p-value less than 0.05. B. Gene set enrichment analysis of genes in AMPK^CA^ livers compared to control livers following 2 months of HFD containing doxycycline. C. Western blot analysis of livers from control and AMPK^CA^ mice following 15 days of HFD containing doxycycline. D. Volcano plot of HNF4α binding sites altered in livers from control and AMPK^CA^ mice following 15 days of HFD containing doxycycline. E. Genomic location and directionality of change in HNF4α binding sites in livers from control and AMPK^CA^ mice following 15 days of HFD containing doxycycline. TTS - Transcriptional termination site. F. Representative HNF4α binding sites in control vs AMPK^CA^ livers. G. Heatmap analysis of HNF4α target genes significantly changed (fdr<0.05) in AMPK^CA^ livers compared to control livers following 2 months of HFD containing doxycycline. Log2 fold change. H. Growth curve of xenograft of MHCC-97H cells expressing wild-type or S304D mutant HNF4α. Inset: representative image of tumor sizes at endpoint n=11. I. Weight of xenografts of MHCC-97H cells expressing wild-type or S304D mutant HNF4α n=11. Welch t-test J. Gene expression analysis of HNF4α targets in xenograft of MHCC-97H cells expressing wild-type or S304D mutant HNF4α. n=11 ±SEM Fisher LSD test *<0.05, **<0.01, ***<0.001, ****<0.0001

Prior studies have shown that AMPK can directly phosphorylate and inhibit HNF4α activity^91^. Consistent with this, we observed a robust increase in phosphorylation of HNF4α at serine 304— a known AMPK phosphorylation site^91^ —in AMPK^CA^ livers under both chow and high-fat diet conditions, compared to control livers (Figures 5C, S5D). We also assessed HNF4α phosphorylation in non-tumor and tumor liver tissue and found strong phosphorylation in non-tumor AMPK^CA^ samples relative to both control tissue and tumors from AMPK^CA^ mice, consistent with the loss of AMPK activity in tumors (Figure S5E). These findings support previous reports that AMPK can directly phosphorylate HNF4α.

Previous studies have shown that AMPK-dependent phosphorylation of HNF4α can inhibit its transcriptional activity by disrupting dimer formation and reducing protein stability^91^. To further clarify how AMPK phosphorylation affects HNF4α’s transcriptional activity, we performed ChIP-Seq analysis of HNF4α in liver samples from mice fed HFD containing doxycycline for 15 days. This time point was selected to capture early regulatory effects of AMPK activation while minimizing confounding effects due to differences in fatty liver development between genotypes. ChIP-Seq was performed on three independent pairs of control and AMPK^CA^ livers. For each HNF4α binding site, we calculated fold change within each pair, p-values, and a resulting z-scores.

Approximately 39,000 peaks were identified with ∼5,235 localized to promoters and transcriptional start sites (Figure 5D). Using a z-score threshold of ±0.75, we identified 788 promoter-associated peaks (-1kb to +100bp) with significantly reduced HNF4α binding across all replicates, compared to only 118 peaks with increased binding (Figure 5E). Similarly, we found decreased binding at 534 exonic peaks and increased binding at only 162 exonic sites in AMPK^CA^ livers. In contrast, we observed increased peak numbers in intronic (1,847 up vs. 1,186 down) and intergenic regions (1,366 up vs. 681 down), and no significant difference in peaks at transcriptional termination sites (52 up vs. 63 down) (Figure S5F). These data suggest that AMPK activation selectively reduces HNF4α binding at promoters and exons.

Genome browser visualization confirmed strong HNF4α binding at the Cyp7a1 promoter in both genotypes, consistent with unchanged expression of this gene (Figures 4I, 5F). However, we observed reduced binding at the promoters of *Abcc3* and *Nr1i3* in AMPK^CA^ livers, consistent with their decreased expression (Figures 4J, 4L, 5F). To investigate whether these changes in HNF4α chromatin binding contribute to transcriptional alterations, we cross-referenced our ChIP-Seq and RNA-Seq datasets. We found that 13% of genes differentially expressed in AMPK^CA^ livers (85 of 647) contained at least one unique HNF4α peak in their promoter or exon regions (Figure 5G). These genes exhibited both up- and down-regulation, reflecting HNF4α’s dual role as a transcriptional activator and repressor depending on context and cofactors^92,93^.

To determine whether AMPK-dependent phosphorylation of HNF4α contributes to liver cancer suppression, we overexpressed either wild-type HNF4α or a phosphomimetic mutant (S304D, where serine 304 is replaced by aspartic acid) in the human HCC cell line MHCC-97H. This mutation mimics constitutive phosphorylation of HNF4α. Following confirmation of transgene expression, cells were injected into the bilateral flanks of SCID immunodeficient mice (Figure S5G). Tumor growth was monitored by caliper measurements, and tumors were weighed at the endpoint. The S304D mutation significantly inhibited tumor growth, resulting in smaller tumors compared to wild-type HNF4α-expressing controls (Figures 5H–I). Expression of the transduced constructs in the tumors was confirmed by western blot and qRT-PCR analyses (Figure S5H–I). To assess whether HNF4α phosphorylation can reproduce the gene expression patterns observed in AMPK^CA^ livers, we analyzed the expression of bile acid biosynthesis and transporter genes in the xenograft tumors. The S304D mutant suppressed expression of *Abcc3*, *Asbt*, and *Nr1i3*, while expression of *Cyp7a1*, *Cyp8a1*, and *Shp* remained unchanged—mirroring the transcriptional profile seen in AMPK^CA^ livers (Figure 5J). These findings support the role of HNF4a phosphorylation in mediating AMPK-dependent transcriptional reprogramming and tumor suppression in the liver.

### Pharmacological AMPK activation ameliorates diabetes and prevents HCC

To confirm the efficacy of AMPK activation in preventing HCC development, we used MK-8722, a potent and selective direct allosteric pan-AMPK activator^94,95^. To minimize stress and better mimic pharmaceutical use in humans—typically administered during the day with food—we delivered MK-8722 via diet at a dose of 10mg/kg^95^. On-target drug activity was confirmed by western blot analysis showing increased phosphorylation of AMPK substrates (P-S79 ACC, P-S792 RAPTOR, P-S555 ULK1, and P-S1387 TSC2) after seven days of HFD feeding (Figure 6A,D), indicating robust AMPK activation.

**Figure 6.**
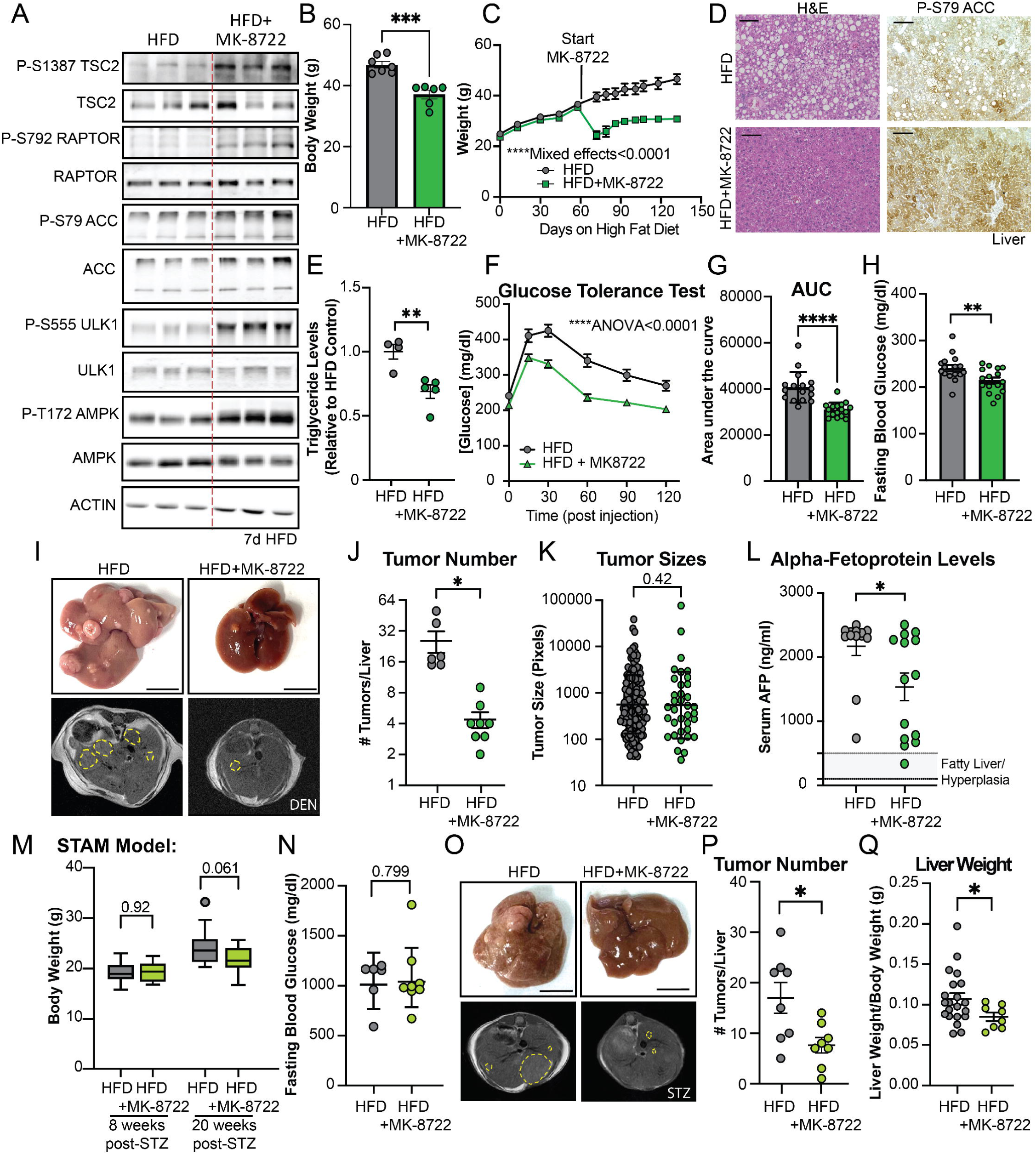
Pharmacological AMPK activation ameliorates diabetes and prevents HCC A. Western blot analysis of female mouse livers fed high fat diet (HFD) containing MK-8722 to activate AMPK for 7 days compared to control HFD livers. B. Body weight (grams) of male mice fed HFD or HFD+MK-8722 for 10 weeks, n=6-7. ±SEM Welch t-test. C. Weight curve of male mice placed on HFD+MK-8722 following 8 weeks on HFD, n=5-8. ±SEM Mixed Effects Analysis. D. H&E analysis of livers from male wild-type mice fed HFD or HFD+MK-8722 for 4 months. Scale bar 100µm. Right: Immunohistochemical analysis of P-ACC in livers from male wild-type mice fed HFD+MK-8722 for 4 months. Scale bar 100µm. E. Quantification of triglyceride levels in liver from male mice on HFD+MK-8722 for 4 months. n=4-5. Welch T-test F. Glucose tolerance test (1mg/ml Glucose IP; left) of male mice on HFD+MK-8722 for 8 weeks. n=16. Two-way ANOVA. G. Area under the curve of GTT analysis in male mice fed HFD or HFD+MK-8722 for 8 weeks. n=16. ±SEM Welch t-test, H. Fasting blood glucose in male mice following 5-6 hour fast. n=16. ±SEM Welch t-test. I. Representative whole mount (top) and MRI (bottom) image of male mice on HFD or HFD+MK-8722 9 months post-DEN injection, Scale bar 1cm. J. Quantification of tumor number from male mice fed HFD or HFD+MK-8722 9-months post-DEN injection by MRI images n=6-8 ±SEM. Welch t-test, K. Quantification of tumor sizes from male mice fed HFD or HFD+MK-8722 9-months post-DEN injection by MRI images n=36-153. Geometric mean +/-SD. Welch t-test, L. Quantification of serum alpha-fetoprotein (AFP) levels in male mice fed HFD or HFD+MK-8722 9-months post-DEN injection. n=13-14. ±SEM Welch t-test. M. Body weight (grams) of male mice fed HFD or HFD+MK-8722 for 8 or 20 weeks post-STZ injection, n=8-20. Fisher LSD test. N. Fasting blood glucose following 5-6 hour fast 20 weeks post-STZ injection n=6-8. Welch t-test. O. Representative whole mount (top) and MRI (bottom) image of male mice on HFD or HFD+MK-8722 20-weeks post-STZ injection, Scale bar 1cm. P. Quantification of tumor number from male mice fed HFD or HFD+MK-8722 20-weeks post-STZ injection by MRI images n=8 ±SEM. Welch t-test. Q. Quantification of liver weight relative to body weight in male mice injected with streptozocin for 20 weeks and fed HFD or HFD+MK-8722. n=8-21. Welch t-test *<0.05, **<0.01, ***<0.001, ****<0.0001

Next, we assessed the drug’s impact on metabolic dysfunction. To model human clinical conditions—where pharmaceuticals are generally introduced after disease onset—we induced obesity and diabetes in wild-type mice through 8 weeks (2 months) of HFD feeding (Figure S6A). Diabetes was confirmed by glucose tolerance testing relative to mice fed chow diet (Figures S6C, D). Mice were then randomized into two groups: one continued on HFD alone, and the other received HFD supplemented with MK-8722. Mice treated with MK-8722 exhibited significant and sustained weight loss with minimal adverse effects (Figures 6B–C, S6E), an effect also observed in female mice (Figure S6F). In contrast, mice on HFD alone continued to gain weight (Figures 6C, S6E–F). This weight loss in mice on MK-8722 correlated with a reduction in hepatic triglyceride levels (Figure 6D–E).

Furthermore, MK-8722 treatment significantly improved glucose tolerance, demonstrating that pharmacological AMPK activation can reverse HFD-induced metabolic disease (Figures 6F–H). Notably, AMPK activation was also observed in peripheral tissues such as epididymal white adipose tissue, suggesting a systemic component to the observed effects (Figure S6G). Importantly, we did not detect adverse cardiac effects, including changes in heart weight, ejection fraction or ejection shortening, which have been reported with this compound at higher doses (Figure S6H–J)^96^.

We next investigated whether pharmacological activation of AMPK could prevent HCC development, similar to our findings in the genetic model. Using the same experimental timeline—8 weeks of HFD followed by treatment with MK-8722 in the HFD—we assessed tumor development in male mice that had been injected with DEN at 2 weeks of age. Tumor burden was evaluated 9 months post-DEN injection using MRI and whole-mount imaging (Figures 6I, S6A). We observed a significant reduction in HCC formation, as indicated by decreased tumor number, tumor size, and serum alpha-fetoprotein levels in MK-8722-treated mice (Figures 6J–L). Of note, histological analysis by H&E staining revealed that tumors from MK-8722-treated mice contained fewer lipid droplets compared to tumors from mice fed HFD alone (Figure S6K). A similar reduction in tumor number was observed in female mice treated with MK-8722 (Figure S6L).

To further evaluate the therapeutic potential of AMPK activation, we tested MK-8722 in the STAM model of diabetes-associated HCC. As previously described, male mice were administered STZ at postnatal day 2 to induce Type 1 Diabetes, followed by HFD from 4 weeks of age to induce fatty liver. At six weeks, when mice in the STAM model exhibit MASLD, they were switched to either HFD alone or HFD containing MK-8722 (Figure S6B)^48^. At eight weeks of age, body weights were similar between groups (Figure 6M). By 20 weeks, we observed a modest reduction in body weight in MK-8722-treated mice, but no improvement in fasting blood glucose levels (Figure 6M– N), consistent with AMPK’s limited impact on glucose homeostasis in the context of insulin deficiency.

Finally, we assessed HCC tumor formation in the STAM model at 20 weeks of age. As in the DEN model, MK-8722 treatment significantly reduced tumor number (Figure 6O–P). This was accompanied by a reduction in liver weight relative to total body weight (Figure 6Q). Together, these findings demonstrate that pharmacological AMPK activation effectively suppresses HCC development, consistent with the tumor-suppressive effects observed in our genetic model.

### Pharmacological AMPK induces bile acid levels and inhibits HNF4a

Because both genetic and pharmacological activation of AMPK inhibited HCC development, we sought to identify a shared mechanism underlying this tumor-suppressive effect. To this end, we performed targeted metabolomic analysis on livers from mice fed HFD followed by HFD alone or HFD supplemented with MK-8722 for 2 months. From this analysis, we identified 188 total metabolites, of which 46 exhibited significant changes (Figure 7A). Surprisingly, when comparing metabolite changes induced by MK-8722 treatment with those observed in AMPK^CA^-expressing livers after 2 months, we found minimal overlap (Figure S7A). The metabolites increased in both contexts included 4-hydroxybutanoic acid, choline, and the bile acids cholic acid (CA) and β-muricholic acid (βMCA). Metabolites consistently downregulated in both models included xanthine and 3-hydroxyanthranilic acid (linked to nucleotide metabolism), methyl pyruvate and 2-methylglutamic acid (byproducts of the TCA cycle), and acetyl-L-carnitine (from fatty acid metabolism). Notably, several TCA cycle intermediates—glutamic acid, fumaric acid, and malic acid—were upregulated following MK-8722 treatment but downregulated in AMPK^CA^ livers (Figures 7A, S4C). These opposing patterns likely reflect the difference between liver-specific AMPK activation and systemic AMPK activation resulting from pharmacologic treatment.

**Figure 7.**
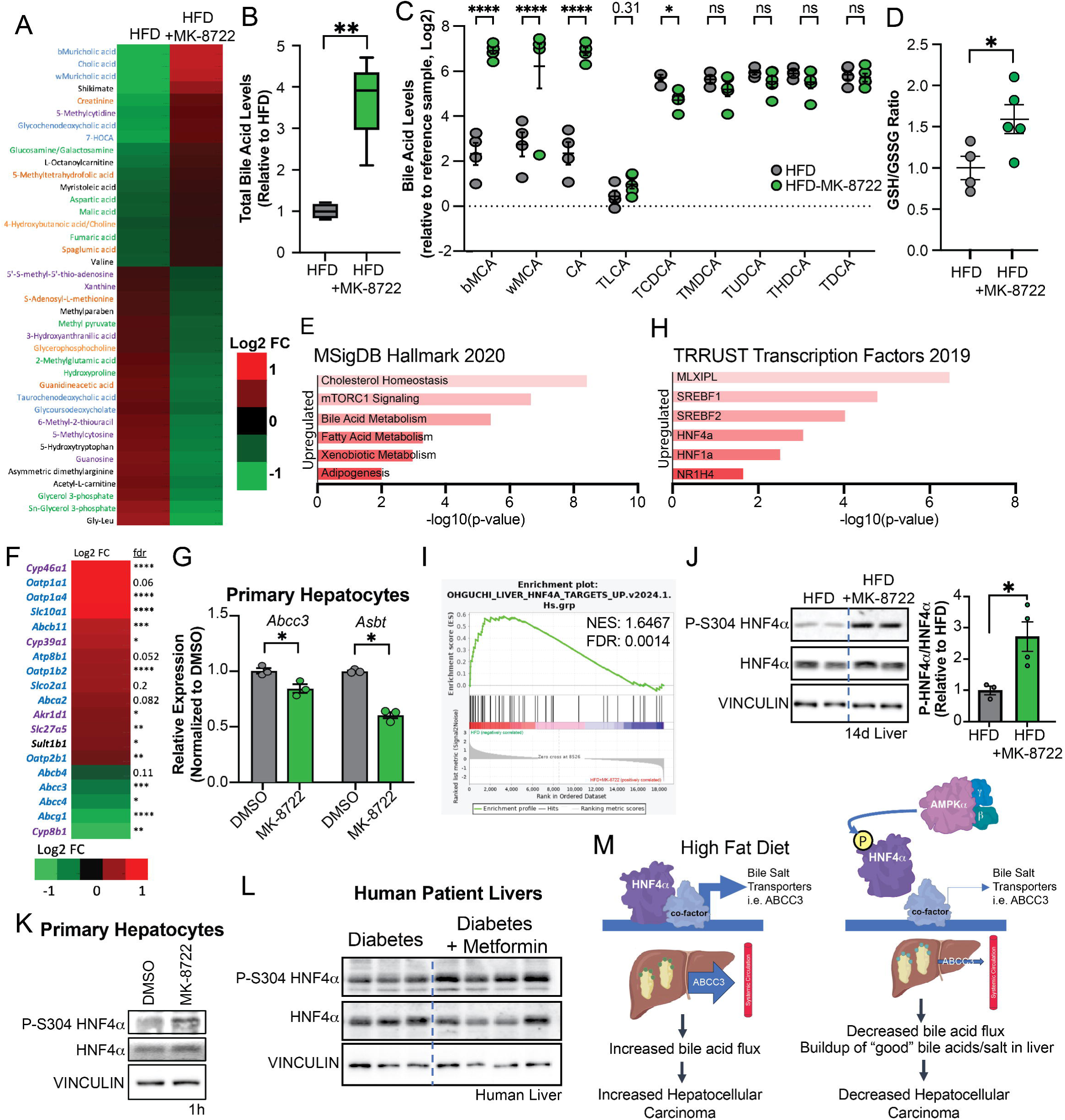
Pharmacological AMPK activation increases bile acid levels and inhibits HNF4aA. A. Heatmap of the log2 fold change for significantly (p-value<0.05, FC>1.5) altered metabolites in liver fed HFD+MK-8722 for 2 months relative to HFD, n=4-6. B. Quantification of total bile acid levels in livers from female mice on HFD versus HFD+MK-8722, n=4-5. Welch t-test, C. Quantification of bile acids levels relative to reference sample, n=4-5. ±SEM. Fisher LSD t-test. D. Relative Glutathione (GSH) to Glutathione disulfide ratio (GSSG) ratio in livers from mice following 2-month treatment with high fat diet containing MK-8722, n=4-5. ±SEM. Welch t-test E. Biological pathways enriched in mice fed HFD+MK-8722 for 2 months relative to control mice fed HFD using Enrichr. F. Heatmap of log2 fold change of gene expression from livers of male mice on HFD+MK-8722 relative to HFD for 2 months, *fdr, Blue: transporters genes, Purple: synthesis genes. G. Relative expression of bile acid transporters in primary mouse hepatocytes treated with DMSO or 10µM MK-8722 for 6 hours, n=3. ±SEM. Welch t-test. H. Transcription factors enriched in mice fed HFD+MK-8722 for 2 months relative to control mice fed HFD using Enrichr. I. Gene set enrichment analysis of genes in mice fed HFD+MK-8722 for 2 months relative to control mice fed HFD. J. Western blot analysis of livers from mice fed HFD+MK-8722 relative to control mice fed HFD for 14 days. Right: Quantification of P-HNF4α levels n=3-4. ±SEM. Welch t-test K. Western blot analysis of primary mouse hepatocytes treated with DMSO or 10µM MK-8722 for 1 hour. L. Western blot analysis of human livers from diabetic individuals treated with or without Metformin. M. Schematic of AMPK’s ability to inhibit HCC development *<0.05, **<0.01, ****<0.0001

The consistent upregulation of bile acids in both models prompted us to quantify total hepatic bile acid levels in mice treated with MK-8722. We observed a significant increase in total bile acid levels, characterized by a low secondary-to-primary bile acid ratio and a low conjugated-to-unconjugated ratio (Figures 7B, S7B–C). Further analysis revealed robust upregulation of primary bile acids, including muricholic acid (MCA) and cholic acid (CA) (Figure 7C). This pattern contrasts with the bile acid profile in AMPK^CA^ mice, where increases were observed in both secondary and conjugated bile acids. However, TLCA levels trended upward in both contexts, suggesting some conserved regulatory effects of AMPK activation. The distinct bile acid species induced by MK-8722 may reflect its action in tissues beyond hepatocytes, including cholangiocytes and intestinal cells. Nonetheless, the bile acids enriched in both contexts are considered less toxic and more hepatoprotective^52,62^. This is indicated by the improved oxidative stress state of the livers from mice treated with MK-8722 as determined by a higher GSH/GSSG ratio (Figure 7D), supporting a model in which AMPK activation fosters a liver environment resistant to injury and tumorigenesis

We next performed RNA-seq on livers from mice fed HFD with or without MK-8722 treatment for 2 months to assess transcriptional changes. We identified 413 genes significantly upregulated, and 911 genes downregulated in response to MK-8722 relative to HFD alone. Enrichment analysis using Enrichr and the MSigDB Hallmark gene sets revealed significant upregulation of pathways related to bile acid metabolism, cholesterol homeostasis (e.g., *Hmgcr*, *Acss2*), mTORC1 signaling (e.g., *Insig1*, *Acly*), and fatty acid metabolism (e.g., *Acaca*, *Fasn*) (Figure 7E). Conversely, downregulated genes were enriched in pathways associated with interferon responses and epithelial-to-mesenchymal transition (Figure S7D).

We next examined expression levels of key bile acid biosynthesis genes^53^. While no changes were observed in *Cyp7a1*, *Cyp27a1*, or *Cyp7b1* (Figure S7E), we found increased expression of alternative pathway enzymes *Cyp46a1* and *Cyp39a1*, along with decreased expression of *Cyp8b1*—a pattern that closely mirrors what we observed in AMPK^CA^ livers (Figure 7F). In parallel, bile acid transporter genes were differentially regulated: expression of hepatic importers such as *Slc10a1* and *Abcb11* was significantly upregulated, while exporters including *Abcc3* and *Abcc4* were downregulated, consistent with enhanced hepatic bile acid retention. To determine whether these transcriptional changes are directly regulated by AMPK activation, we treated primary hepatocytes from wild-type mice with MK-8722 for six hours. This resulted in robust downregulation of *Abcc3* and *Asbt* expression compared to DMSO controls (Figure 7G), supporting a direct role for AMPK in controlling bile acid transporter gene expression.

Importantly, although global gene expression analysis revealed only ∼10% overlap in differentially expressed genes between MK-8722-treated livers and previously published AMPK^CA^ datasets (Figures S7F–G), many of the shared genes are involved in bile acid metabolism and transport— underscoring a convergent molecular signature across both pharmacological and genetic models. Furthermore, Hallmark pathway analysis demonstrated nearly complete overlap in enriched biological processes, including bile acid metabolism, cholesterol homeostasis, and fatty acid biosynthesis (Figures 7E, S5A). This suggests that despite modest gene-level overlap, both interventions engage in a common regulatory program through AMPK activation to mitigate hepatocellular carcinoma risk.

To further define regulatory mechanisms, we used Enrichr with the TRRUST database to identify transcription factors enriched among genes upregulated by MK-8722. The top hits included key regulators of cholesterol and fatty acid biosynthesis such as MLXIPL and SREBP1/2, as well as HNF4α (Figure 7H). In contrast, transcription factors enriched among downregulated genes were predominantly immune-related, including STAT1, RELA, and NFKB1 (Figure S7H). Gene set enrichment analysis further highlighted HNF4α as a significantly modulated factor (Figure 7I, S7I). We therefore examined phosphorylation of HNF4α at serine 304 in liver tissue and primary hepatocytes treated with MK-8722. In both contexts, we observed increased phosphorylation of HNF4α relative to HFD controls (Figure 7J–K), consistent with AMPK-dependent inhibition of HNF4α activity as previously observed in our genetic models.

To directly assess HNF4α phosphorylation in human patient samples, we analyzed liver tissue from individuals with diabetes treated with or without metformin, a known AMPK activator. We observed increased levels of phosphorylated HNF4α in the liver tissue of metformin-treated patients, indicating that AMPK activation is sufficient to induce HNF4α phosphorylation in human liver (Figure 7L). These results parallel our findings in genetic mouse models, demonstrating that pharmacological AMPK activation suppresses HNF4α activity through the same phosphorylation-dependent mechanism. Thus, AMPK drives HNF4α phosphorylation in both mouse and human liver tissues, particularly in contexts where HCC is known to be prevented^6–12^, underscoring a unified, targetable pathway for HCC prevention.Together, these findings support a model in which AMPK activation suppresses HCC by phosphorylating and inhibiting HNF4α (Figure 7M). This mechanism reduces bile acid-driven hepatotoxicity and mitigates transcriptional programs associated with tumorigenesis.

## Discussion

Our study demonstrates that AMPK activation significantly reduces tumor formation in both DEN-induced and STAM models of HCC. This finding underscores the potential of AMPK activation as a potent strategy for preventing HCC development, particularly in the context of metabolic disorders such as obesity and Type 2 Diabetes. The mechanistic insights provided by our study reveal that AMPK activation influences bile acid metabolism and HNF4α signaling, which are crucial for HCC prevention.

The role of AMPK in cancer is complex and context-dependent, with evidence supporting both tumor-suppressive and tumor-promoting functions. In several malignancies, including breast, prostate, and melanoma, AMPK activation has been associated with reduced tumor growth, often through mechanisms such as inhibition of de novo lipogenesis, suppression of mTOR signaling, and induction of autophagy or apoptosis^97,98^. For instance, in prostate cancer, both pharmacologic and genetic activation of AMPK suppressed tumor development by promoting mitochondrial biogenesis and fatty acid oxidation while downregulating inflammatory and proliferative pathways^99,100^. However, emerging studies have also highlighted tumor-promoting roles for AMPK in specific contexts. In lung cancer, Myc-induced lymphoma, and acute myeloid leukemia (AML), AMPK loss paradoxically impaired tumor development, likely due to metabolic vulnerabilities such as lysosomal dysfunction or energetic stress^44,101–103^. These findings underscore the tissue-specific and context-dependent nature of AMPK signaling in cancer. In HCC, preclinical studies have shown that AMPK activation, including through agents like metformin, can suppress HCC cell growth by inhibiting glycolysis and promoting metabolic reprogramming^12,104,105^. Despite these insights, the definitive role of AMPK in HCC tumorigenesis had remained unresolved, necessitating our *in vivo* studies using genetically engineered mouse models to clarify its function in liver cancer development.

In our genetic model of constitutive AMPK activation, we made the striking observation that AMPK was consistently downregulated at both the activity and protein levels in tumor tissues compared to matched non-tumor tissue. This finding aligns with previous report linking low AMPK expression to poor prognosis in HCC cancer patients^106^. The mechanism underlying this downregulation remains unclear but may involve impaired translation or reduced protein stability. One possibility is that prolonged AMPK activation suppresses its own translation via feedback inhibition of mTORC1, a central regulator of protein synthesis^18^. Alternatively, tumor cells may actively degrade AMPK protein through ubiquitin-mediated pathways. For instance, SAPS3, a regulatory subunit of protein phosphatase 6, has been shown to inhibit AMPK by promoting its dephosphorylation, particularly under metabolic stress conditions such as high-fat diet exposure^107^. Additionally, cancer-specific E3 ubiquitin ligases such as MAGE-A3/6 and TRIM-72 have been implicated in targeting AMPK for degradation, with TRIM-72 activity enhanced by AKT signaling^108,109^. These observations suggest that tumors can evolve strategies to bypass the tumor-suppressive effects of AMPK, potentially even in our context of constitutive activation. This represents an intriguing avenue for future investigation, with potential implications for understanding resistance to metabolic therapies targeting AMPK.

One of the key findings of our study is that AMPK activation increases hepatic bile acid levels, particularly taurine-conjugated bile acids, which are generally considered less cytotoxic and more hepatoprotective. This protective effect is supported by improved liver oxidative status in both MK-8722-treated and AMPK^CA^ mice on high fat diet. However, the role of bile acids in HCC remains complex and, at times, contradictory. While bile acids are essential for lipid digestion, nutrient absorption, and metabolic regulation, their dysregulation has been linked to liver injury and carcinogenesis^53,59,110,111^. Supporting this duality, agents like UDCA, taurine, and TLCA exhibit anti-inflammatory and hepatoprotective effects, whereas others, such as CDCA, can promote inflammation and pro-growth signaling^62,63,112–115^. Notably, clinical studies have proposed that elevated serum bile acid levels may serve as prognostic markers for HCC^57^. In contrast, analyses of hepatic tissues often show reduced bile acid levels in tumors compared to matched non-tumor samples^116^. Our findings align with this discrepancy: we observed increased hepatic bile acids without corresponding changes in serum levels. This suggests that hepatic and circulating bile acid pools may be regulated independently, reinforcing the potential of targeting hepatic bile acid metabolism for HCC prevention. It is also important to consider sex-based differences in bile acid regulation. Our studies were conducted primarily in male mice, whereas the publicly available RNA-seq datasets were derived from females^45^. Given that genes such as *Cyp39a1* and *Cyp3a11* show sex-dependent expression, this may influence interpretation of gene expression and metabolic outcomes^117,118^. Future studies should account for sex as a biological variable to better understand the regulatory mechanisms linking bile acid metabolism and HCC risk. Overall, our findings reveal a context in which bile acids play a protective role, contributing to the understanding of their function in HCC, and highlighting the intricate and context-dependent nature of their regulation in liver health and disease.

Our study also highlights the critical role of AMPK-HNF4α interaction in HCC development. We found that AMPK activation inhibits HNF4α activity through phosphorylation, which reduces bile acid-driven hepatotoxicity and suppresses transcriptional programs associated with tumorigenesis. The role of HNF4α in HCC development is complex and remains an area of active investigation. For example, HNF4α exists in multiple isoforms, primarily driven by two promoters: P1 and P2^119^. Notably, the P2-driven isoform is upregulated in fatty liver disease and HCC and has been associated with pro-tumorigenic activity, while the P1-driven isoform is considered tumor-suppressive^120–123^. Moreover, HNF4α function is shaped by its interacting partners, which vary by isoform and cellular context^92^. For example, HNF4α can interact with STAT3 and β-catenin to promote oncogenic gene expression, while its interaction with HNF1α supports hepatocyte differentiation and liver function, potentially counteracting tumorigenesis^124–126^. In our study, we observed selective downregulation of HNF4α binding and HNF4α-dependent gene expression following AMPK activation, suggesting that phosphorylation may alter its DNA-binding affinity or co-factor interactions. Interestingly, canonical HNF4α targets such as *Cyp7a1* were not among the AMPK-sensitive binding sites^127,128^. Instead, we identified altered regulation of nuclear receptors like CAR (*Nr1i3*), indicating a possible secondary mechanism by which AMPK modulates gene expression^129,130^. Although previous studies have reported that AMPK can reduce HNF4α protein stability, we did not observe consistent changes in total HNF4α protein levels^91^. This raises the possibility that AMPK may instead influence isoform expression, post-translational modifications, or protein-protein interactions. Future studies should explore whether AMPK alters the ratio of HNF4α isoforms or modulates its interaction network to regulate gene expression and suppress HCC. Regardless, these findings support the idea that targeting HNF4α activity may offer a promising therapeutic strategy in AMPK-driven HCC prevention.

The broader implications of our study emphasize the potential of novel pharmaceutical AMPK activators in HCC prevention and treatment. Both genetic and pharmacological activation of AMPK effectively suppress HCC development, suggesting that AMPK-based therapies could be beneficial for patients with metabolic disorders and those at elevated risk of developing HCC. The development of AMPK activators as therapeutic agents could provide a new avenue for cancer prevention and treatment, particularly in the context of metabolic dysregulation.

In conclusion, our study provides significant insights into the role of AMPK activation in HCC prevention. These findings highlight the importance of bile acid metabolism and HNF4α signaling in mediating the tumor-suppressive effects of AMPK. The overall implications of our study suggest that AMPK activation could be a promising strategy for preventing HCC. Further research is needed to fully understand the mechanisms involved, including HNF4α inhibition, and to develop effective AMPK-based therapies for HCC prevention and treatment.

## Resource Availability

RNA-seq, ChIP-Seq, and metabolomics data will be made publicly available upon publication.

## Acknowledgements

We would like to thank Nicholas Egan and Lillian Eichner for their editorial comments and Monali Naik and Naghmana Ashraf for in-depth discussions. We thank Ruth Wooten-Kee and Bingning Dong for primer sequences and insight into bile acid biology. We thank Dr. Reuben Shaw for the AMPK^CA^ mouse model.

## Author Contributions

Z.S. and B.L. designed and conducted the experiments, analyzed results, and edited the manuscript. C.U. and F.A. performed experiments. N.P. performed metabolomic analyses. J.V.N. designed and performed experiments, analyzed results, and wrote the manuscript.

## Footnotes Funding

This work was supported in part by the Cancer Prevention and Research Institute of Texas (RR210013) to J.L.V.N. J.L.V.N. is a CPRIT Scholar in Cancer Research. This project was supported by the Mouse Metabolism and Phenotyping Core at Baylor College of Medicine which receives funding from NIH (UM1HG006348, R01DK114356) and additional support from the Baylor College of Medicine Advanced Technology Cores and an S10 grant (S10 OD032380). This project was supported by CPRIT Proteomics and Metabolomics Core Facility (N.P.), (RP210227), NIH (P30 CA125123), and Dan L. Duncan Cancer Center.

## Materials and methods

### Mouse Studies

AMPK^CA^ mice were previously described^45^. In summary, C57Bl/6J mice harboring a constitutive-active truncation mutant of AMPKa1 (amino acids 1-312) downstream of a tetracycline-responsive promoter in the Col1A1 locus. The mice also harbor a Cre-inducible rtTA3 cassette in the Rosa26 locus. These mice were crossed to Albumin-Cre mice (Jackson Laboratories #003574). To induce HCC, a single dose of 25 mg/kg DEN (N-Nitrosodiethylamine, Sigma-Aldrich, N0756-10ML) was injected intraperitoneally into 14 days old mice^131^. Continuous high fat diet was started around 8 weeks post DEN injection. Alternatively, mice were injected with a single dose of 200mg Streptozocin (STZ) dorsally at 2 days postnatal^48^. Continuous high fat diet was started at 4 weeks of age. For non-tumor studies, mice were 8-10 weeks of age at the beginning of experiments. Males were used for experiments except where noted. Mice were housed with a standard 12-hour day and night cycles. All mice were housed in AAALAC-accredited, specific-pathogen-free animal care facilities at Baylor College of Medicine, and all procedures were approved by the BCM Institutional Animal Care and Use Committee.

### Special Diets

For diet-induced obesity studies, mice were fed a high fat diet (45% calories from fat; D12451), HFD with 87mg/kg doxycycline (*D21060308)* or calorie-matched Chow Diet with 50mg/kg doxycycline (10% kcal fat *D23082904)*^45^. For AMPK activation studies, mice were fed a high fat diet (D12451) containing 100mg/kg MK-8722 (MedChem Express, HY-111363). Assuming mice eat 10% of their body weight each day, this concentration corresponds to a dose 10mg/kg of body weight. Mice on doxycycline-containing diets were housed in the same cage, while mice on diet-containing MK-8722 were housed in cages separate from their control counterparts on HFD.

### MRI Imaging and Echocardiogram

Tumors were imaged by magnetic resonance imaging (MRI) of live mice. Mice were anesthetized with continual flow isoflurane. Both axial and coronal series were collected based on the T2 sequence on the pharmascan with some adjustments. ROI started about 1-2 mm rostral to the liver and then continued caudally for 25 slices – several of the animals would present with extremely enlarged livers. 1mM thickness images were collected with a 256×256 matrix and 3cm field of view. Fat suppression was off and respiratory gated (4x average). TE: 21.62 TR:4000. Images were analyzed by image J and areas and number of tumors quantified. Tumor spanning multiple sections were counted as a single tumor and results verified by comparison with whole-mount images at time of dissection.

For assessment of echocardiogram (ECG), animals were anesthetized using isoflurane anesthesia delivered in oxygen with a precision vaporizer (3% isoflurane induction, 1.5% isoflurane for maintenance during recording). Animals were placed onto a Rodent Surgical Monitor (Indus) in a supine position and paws placed on surface electrodes using conductive paste and secured with surgical tape. Internal body temperature was confirmed via rectal thermometer, and animals were stabilized at 36 – 37.5 C before initiating data collection. The ECG signal was transferred via low latency analog output to a powerlab analog to digital signal acquisition system with LabChart 8 software (AD Instruments). The ECG analysis module of the LabChart software was configured for mouse heartbeat detection and ECG outcome parameters were collected for the left ventricle. Following data collection animals were returned to their home cage and provided with thermal support until sternally recumbent and resumption of species typical behavior.

### Glucose Tolerance (GTT)

Mice were fasted in paper bedding for 6 hours (8:00 am to 2:00 pm) before measuring fasting blood glucose and beginning the tolerance tests. Glucose (1.5 g/kg body weight) was injected intraperitoneally. Blood glucose level was measured using Aviva ACCU-CHEK® glucose meter prior to injection and after injection, at the times indicated. For fasting blood glucose of STZ-injected mice, blood was collected at time of euthanasia following a 4 hour fast and glucose levels measured using a glucose colorimetric assay kit (Raybiotech MAGLU1).

### Tissue Collection

Mice were fed ad-lib or fasted for 4 hours (for afternoon collections) prior to sacrifice through Avertin anaesthetization (Sigma T48402). Mice were collected at either 10am or 2pm. 100 mg/kg BrdU (sigma B5002) was I.P. injected 4h prior to sacrifice. Blood serum was collected via retro-orbital bleeding and serum was obtained by centrifugation. Heart and brain were excised and weighed. Tumor and non-tumor tissues for metabolomic and transcriptional analysis were macroscopically dissected at time of injection and snap frozen in liquid nitrogen. Tissues for immunohistochemistry were formalin fixed overnight and tissues for oil red O were frozen in OCT prior to cryosectioning.

### Cell culture & Xenograft

Human HCC cell line MHCC-97H was obtained from Dr. David Moore. All cell lines were incubated at 37°C and were maintained in an atmosphere containing 5% CO_2_. Cells were grown in Dulbecco’s modified Eagles medium (DMEM) plus 10% fetal bovine serum. Cell identity was confirmed by STR analysis and mycoplasma testing was routinely performed. Cells were infected with lenti-viral constructs containing wild-type HNF4α or a mutated S304D version of HNF4α. HNF4α was cloned from pLV-HNF4A-FOXA2-FOXA3 constructs from addgene (#149724) into pLenti-CMV/TO-Puro empty (addgene 17482) using primers: F-5’GGTACCGAGCTCAGATCTGCCACCATGCGACTCTCCAAAACC and R-5’TTCTTCTCTAGACTAGATAACTTCCTGCTTGGTGATG. The serine 304 site was mutated to aspartic acid using primers: F-5’GCGGCTGCGTgatCAGGTGCAGG and R-5’TTGATCTTCCCTGGATCGC and sequence confirmed by whole-plasmid sequencing. Lenti virus was produced in 293T cell and used to transduce MHCC-97H followed by selection with puromycin. Male Fox Chase SCID Beige Mice (Charles River) (∼8 weeks of age) were used for xenograft studies. Mice were inoculated bilaterally by subcutaneous injection with 4 × 10^6^ cells resuspended in Matrigel Basement Membrane, 1:1 mixture solution. Tumors were allowed to develop to a volume of around 1000 mm^3^. Tumor size was measured biweekly using calipers until collection. Volume was calculated using the formula: (shortest-length/2)^2^ x Longest-Length. Expression of wild-type and mutant HNF4α was confirmed by western blot and qRT-PCR for HNF4a overexpression and by sequencing PCR fragments obtained from cDNA.

### Primary Hepatocytes

Primary mouse hepatocytes were isolated as described previously (Van Nostrand, 2020) from 8- to 16-week-old mice by portal vein perfusion with collagenase (Sigma, C5138). Hepatocytes were plated at 7×10^5^ cells per well of a 6 well plate in DMEM (Cellgro, MT 10-017-CV) containing 5% FBS and Pen/Strep, and allowed to attach fully prior to indicated treatments.

### Immunohistochemistry and Image Analysis

Fixed tissue was processed and paraffin embedded. 5 μm sections were prepared and either stained with eosin and hematoxylin or used for immunohistochemistry. Immunohistochemistry was performed by antigen retrieval using Trilogy (Cell Marque; Sigma 920P). Slides were boiled for 15 minutes in Trilogy, allowed to cool, and blocked with ImmPRESS IHC kit (Mouse or Rabbit; Vector Labs MP740215, MP740115). Staining was developed using DAB kit (VectorLabs SK4105) and counter-stained with Harris hematoxylin. Slides were imaged using a Keyence microscope and image analysis was performed using Keyence analysis software. For quantification, when possible, multiple 10x images at the same exposure were taken across the tissue of interest and the resulting quantification averaged to get a single value per sample. The area was calculated as the total area minus the area of lipid droplets to account for differences in cell number per image. Pixel intensity was determined per image and normalized to the calculated area. For Oil Red O, samples were embedded in OCT and cryosectioned. Samples were stained with 1.8mg/ml Oil Red O and modified Harris hematoxylin, according to standard protocols^104^. Samples were blindly scored for levels of staining, using criteria including lipid size and lipid number.

### Protein Extraction and Immunoblotting, Western Blotting

Upon euthanization, tissue and tumors were harvested, immediately snap frozen in liquid nitrogen and homogenized on ice in CST lysing buffer (20 mM Tris-HCl pH 7.5, 150 mM NaCl, 1 mM EDTA, 1 mM EGTA, 0.1% SDS, 50 mM sodium fluoride, 2.5 mM sodium pyrophosphate, 2 mM beta-glycerophosphate, 1 mM Na3VO4, 10 nM Calyculin A) supplemented with protease inhibitors (Roche, cOmplete™, #11836170001). Primary hepatocytes were lysed on ice for 10 min with shaking in lysis buffer. Lysates were centrifuged at 16,000 x g for 10 min at 4°C. Protein concentration was determined using Pierce™ BCA protein assay kit (Thermo Scientific, #23225). Lysates were resolved on 8%–12% SDS-PAGE gels, depending on the experiment, and transferred to PVDF membranes for immunoblotting. Membranes were blocked in 5% nonfat dry milk or 5% BSA in Tris buffered saline containing 0.1% Tween 20 for 1 h and probed with various antibodies shown below. Immunoreactive bands were detected by SuperSignal™ West Femto Maximum Sensitivity Substrate (Thermo Scientific™, 34096) and revealed by azure biosystems 300. When appropriate, protein levels were quantified by densitometry analysis using ImageJ.

### Tissue and Serum Analyses

For alpha-fetoprotein quantification, serum was measured by mouse a-Fetoprotein Quantikine ELISA Kit (R&D Systems, MAFP00). TBARs in the liver were quantified per the manufacturer’s instructions using the TBARS (TCA Method) Assay Kit (Cayman,700870). TBARs were normalized to protein amount determined by BCA method.

### Bile Acid, Triglyceride, and Metabolite analysis

Lipid extraction, mass spectrometry (MS) data acquisition, MS raw data processing, and analysis were described previously in detail^132^. Metabolomics analyses were performed on liver samples collected at various timepoints by the Metabolomics Core at Baylor College of Medicine as described previously^49^. Snap frozen liver and serum were submitted to the Metabolomics Core at Baylor College of Medicine for relative bile acid measurement by HPLC as previously described^133,134^. Data was acquired with Agilent mass hunter acquisition software and analyzed with Mass hunter workstation quantitative software. 4-6 tissue samples were used for each genotype or treatment. For metabolic analyses, a p-value of <0.05 and a fold change above 1.5 relative to control sample at each timepoint was used to identify metabolites of interest.

### RNA Isolation, Quantitative Real-Time PCR, and RNA-Seq

RNA was isolated from liver, ileum of the intestine, primary hepatocytes, and tumor tissue using RNeasy tissue mini kit (QIAGEN cat # 74804) or Trizol/Vezol extraction, and cDNA synthesis performed using SuperScript III First-Strand synthesis system (Invitrogen cat # 18080051) or cDNA Synthesis Kit (New England Biolabs). Expression levels were determined by quantitative PCR (Taq Pro Universal SYBR qPCR Master Mix, Vazyme, Q712:03) on the Real-Time PCR System (ViiA7 by lift technologies, AB applied biosystems). The relative expression of target genes was normalized by β-Actin expression as an internal control.

For RNA-seq, RNA integrity (RIN) numbers were determined using the Agilent TapeStation prior to library preparation. Messenger RNA was purified from total RNA using poly-T oligo-attached magnetic beads. After fragmentation, the first strand cDNA was synthesized using random hexamer primers, followed by the second strand cDNA synthesis. The library was checked with Qubit and real-time PCR for quantification and bioanalyzer for size distribution detection. Quantified libraries were pooled and sequenced on Illumina NovaSeq PE150 platform at Novogene. Raw sequencing data were processed by removing reads containing adapter, reads containing ploy-N and low-quality reads from raw data. Reads were mapped to reference genome using Hisat2 v2.0.5. featureCounts[3] v1.5.0-p3 was used to count the reads numbers mapped to each gene, and FPKM of each gene was calculated based on the length of the gene and reads count mapped to this gene. Differential expression analysis was performed using the DESeq2Rpackage (1.20.0). The resulting p-values were adjusted using the Benjamini and Hochberg’s approach for controlling the false discovery rate. Genes with an adjusted p-value<=0.05 found by DESeq2 and absolute fold change of 1.3 were assigned as differentially expressed. Gene Set Enrichment analysis was performed using Genepattern and enrichment analysis was performed using Enrichr.

Published RNA-seq was obtained from GEO (GSE122767). Gene expression from livers of AMPK^CA^ mice was compared to controls, including wildtype mice treated with and without doxycycline and albumin-cre expressing mice in the absence of doxycycline, with 3 samples per condition. Genes with an adjusted p-value<=0.05 and absolute fold change of 1.3 were assigned as differentially expressed and used for further analysis. Gene Set Enrichment analysis was performed using Genepattern and enrichment analysis was performed using Enrichr.

### ChIP-Seq

Frozen tissue was sent to Active Motif Services (Carlsbad, CA) for ChIP-Seq. Active Motif prepared chromatin, performed ChIP reactions, generated libraries, sequenced the libraries, and performed data analysis. In brief, tissue was submersed in PBS + 1% formaldehyde, cut into small pieces, and incubated at room temperature for 15 minutes. Fixation was stopped by the addition of 0.125 M glycine (final). Chromatin was isolated by adding lysis buffer, followed by disruption with a Dounce homogenizer, washing and sonication using the PIXUL® Multi-Sample Sonicator (Active Motif, Catalog #53130) to shear DNA to an average fragment size of 200–1000 bp. To determine chromatin yield, an aliquot of the sheared chromatin was reverse crosslinked at 65°C, treated with RNase and proteinase K, and subjected to DNA extraction using SPRI beads (Beckman Coulter). DNA concentrations were measured using a Qubit Fluorometer (Thermo Fisher), and total chromatin yield was extrapolated based on the original chromatin volume.

For ChIP reactions, aliquots of chromatin were precleared with protein G agarose beads (Invitrogen). Immunoprecipitations were performed using antibodies specific to the target proteins of interest. After washing, immune complexes were eluted from the beads using SDS buffer, treated with Rnase and proteinase K, and de-crosslinked by overnight incubation at 65°C. ChIP DNA was then purified using phenol-chloroform extraction and ethanol precipitation.

ChIP DNA libraries were prepared using either the PrepX DNA Library Kit (Takara Bio) on the Apollo automation platform or the NEB DNA Library Prep Kit, following the manufacturers’ protocols. Libraries were sequenced on the Illumina platform and the resulting data were analyzed using standard ChIP-Seq workflows. In brief, reads were extended in silico (using Active Motif software) at their 3’-ends to a length of 200 bp, which corresponds to the average fragment length in the size-selected library. To identify the density of fragments (extended tags) along the genome, the genome was divided into 32-nt bins and the number of fragments in each bin was determined. Peaks were called using MACS/MACS2 (Zhang et al., Genome Biology 2008, 9: R137) and SICER (Zang et al., Bioinformatics 25, 1952-1958, 2009). To normalize, the tag number of all samples (within a comparison group) was reduced by random sampling to the number of tags present in the smallest sample. To compare peak metrics, overlapping Intervals were grouped into “Merged Regions,” which were defined by the start coordinate of the most upstream Interval and the end coordinate of the most downstream Interval. After defining the Intervals and Merged

Regions, their genomic locations along with their proximities to gene annotations and other genomic features were determined and average and peak fragment densities within Intervals and Merged Regions were compiled. Fold change of peaks within each experimental set was compared and a p-value generated. Representative UCSC genome browser tracks are shown.

#### Patient Liver Samples

Deidentified liver samples from individuals with diabetes treated with or without Metformin were obtained from the Baylor College of Medicine Human Tissue Acquisition and Pathology Core.

| Antibody | Company | REF |
| --- | --- | --- |
| HNF4A Monoclonal Antibody (K9218) | Invitrogen | MA1-199 |
| Phospho-HNF4A (Ser313) Antibody (aka S304) | Invitrogen | PA5-36857 |
| AMPKalpha Rabbit Antibody | Cell Signaling Technology | 2535L |
| P-AMPKalpha (T172) (40H9) Rabbit mAb | Cell Signaling Technology | 2535L |
| Tuberin/TSC2 (D93F12) XP ® Rabbit mAb | Cell Signaling Technology | 4308S |
| P-Tuberin/TSC2 (S1387) (D2R3A) Rabbit mAb | Cell Signaling Technology | 23402 |
| P-ULK1 (S555) (D1H4) Rabbit mAb | Cell Signaling Technology | 5869 |
| ULK1 | Cell Signaling Technology | 8054S |
| Acetyl-CoA Carboxylase Rabbit mAb | Cell Signaling Technology | 3662S |
| Phospho-ACC (Ser79) Rabbit mAb | Cell Signaling Technology | 11818S |
| RAPTOR (24C12) Rabbit mAb | Cell Signaling Technology | 2280S |
| Phospho-RAPTOR (S792) Rabbit Ab | Cell Signaling Technology | 2083S |
| CSAD | Invitrogen | PA5-120240 |
| VINCULIN (E1E9V) XP Rabbit mAb | Cell Signaling Technology | 13901S |
| β-ACTIN | Sigma | A5441 |
| BrdU | Abcam | ab6326 |
| HNF4α (for ChIP-seq analysis) | Abcam | ab41898 |
| GFP | Cell Signaling Technology | 2956 |
| Phospho-HNF4A (Ser313) Antibody (aka S304) | Abcam | ab78356 |

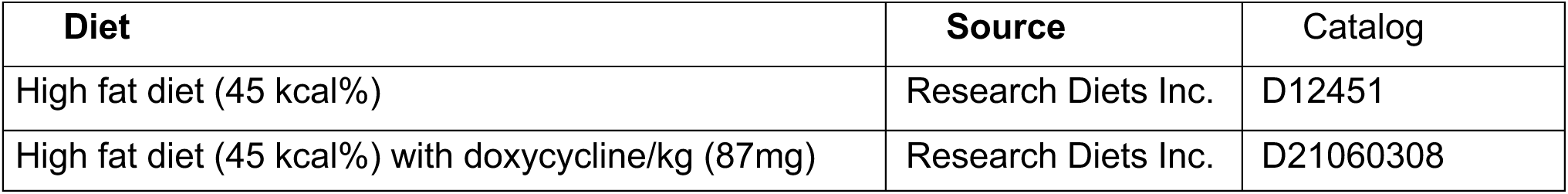
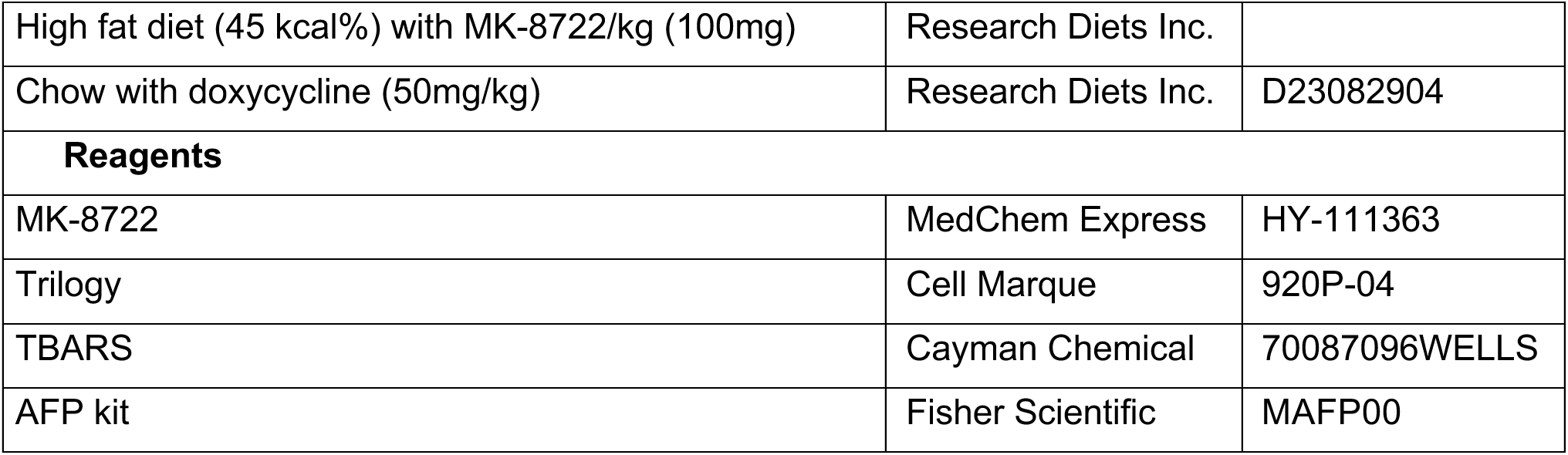

| Diet | Source | Catalog |
| --- | --- | --- |
| High fat diet (45 kcal%) | Research Diets Inc. | D12451 |
| High fat diet (45 kcal%) with doxycycline/kg (87mg) | Research Diets Inc. | D21060308 |
| High fat diet (45 kcal%) with MK-8722/kg (100mg) | Research Diets Inc. |  |
| Chow with doxycycline (50mg/kg) | Research Diets Inc. | D23082904 |
| **Reagents** |  |  |
| MK-8722 | MedChem Express | HY-111363 |
| Trilogy | Cell Marque | 920P-04 |
| TBARS | Cayman Chemical | 70087096WELLS |
| AFP kit | Fisher Scientific | MAFP00 |

#### Oligonucleotides

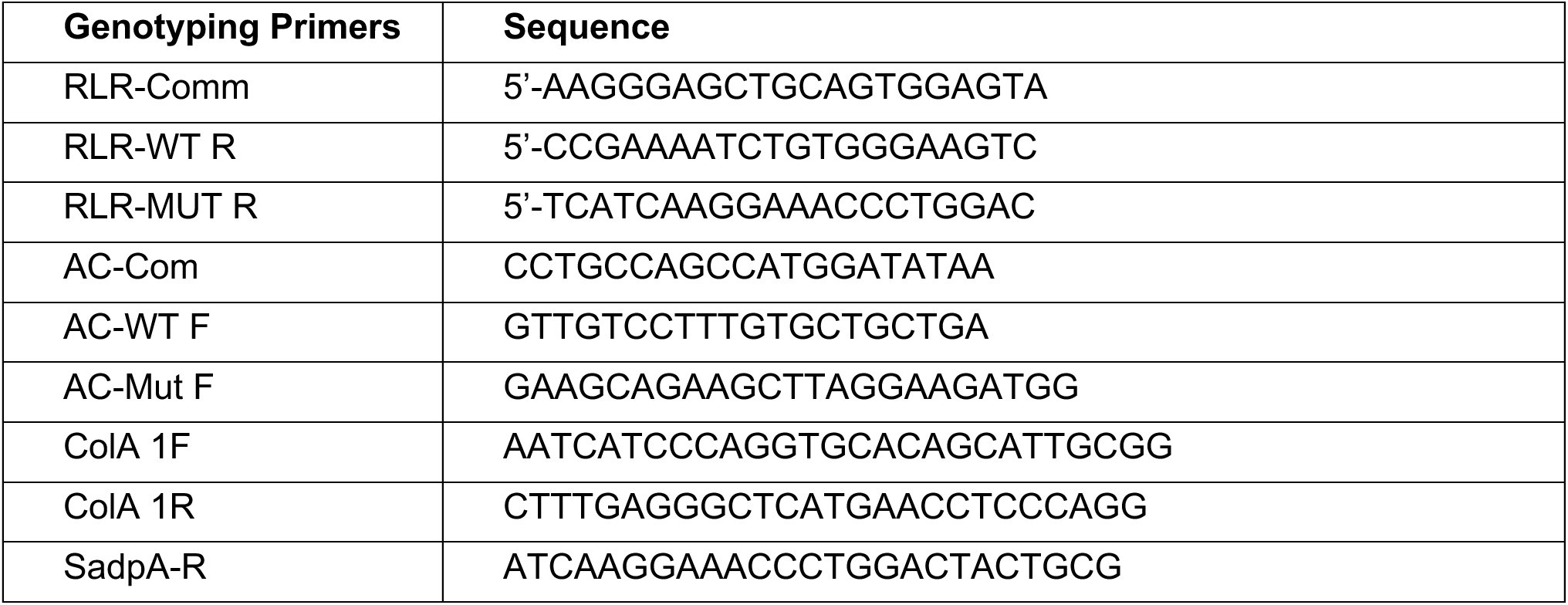

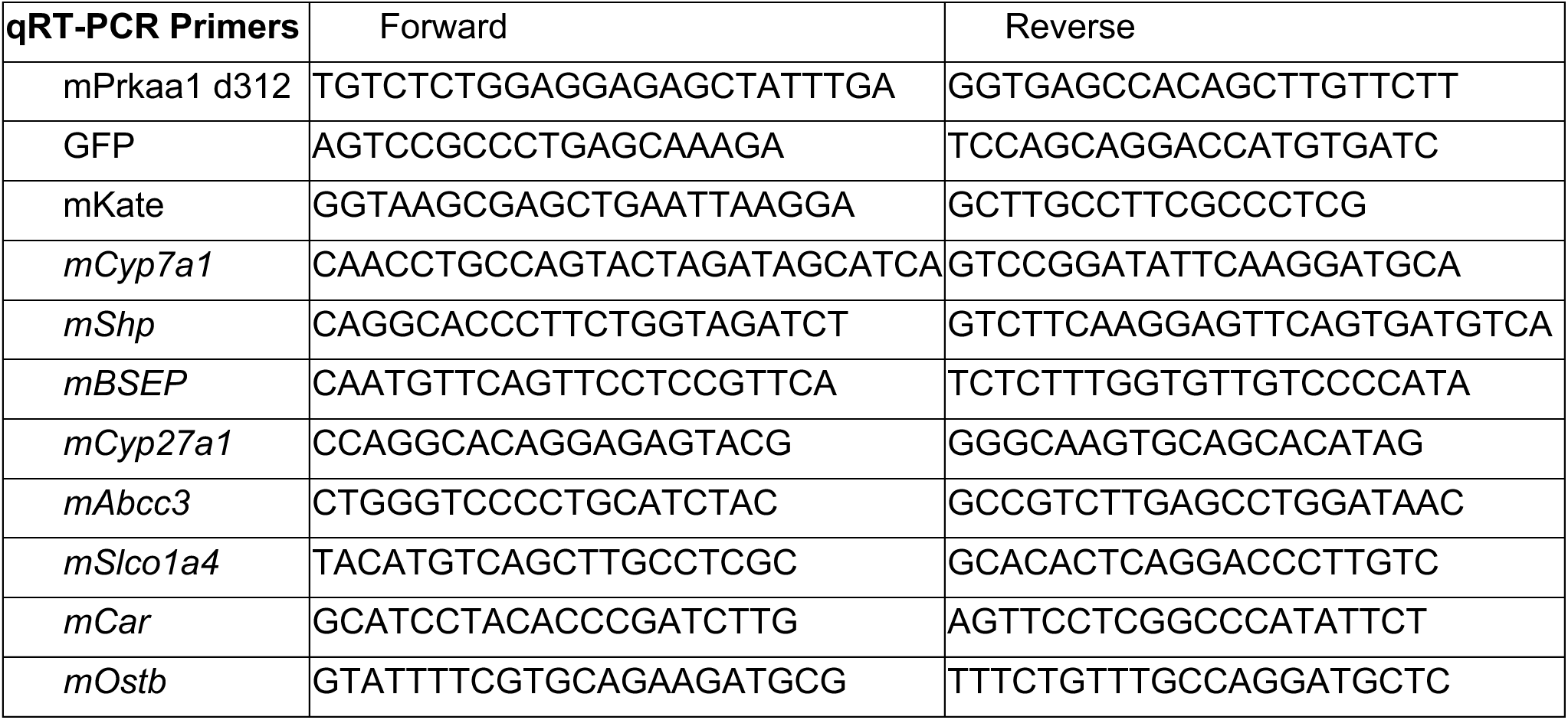
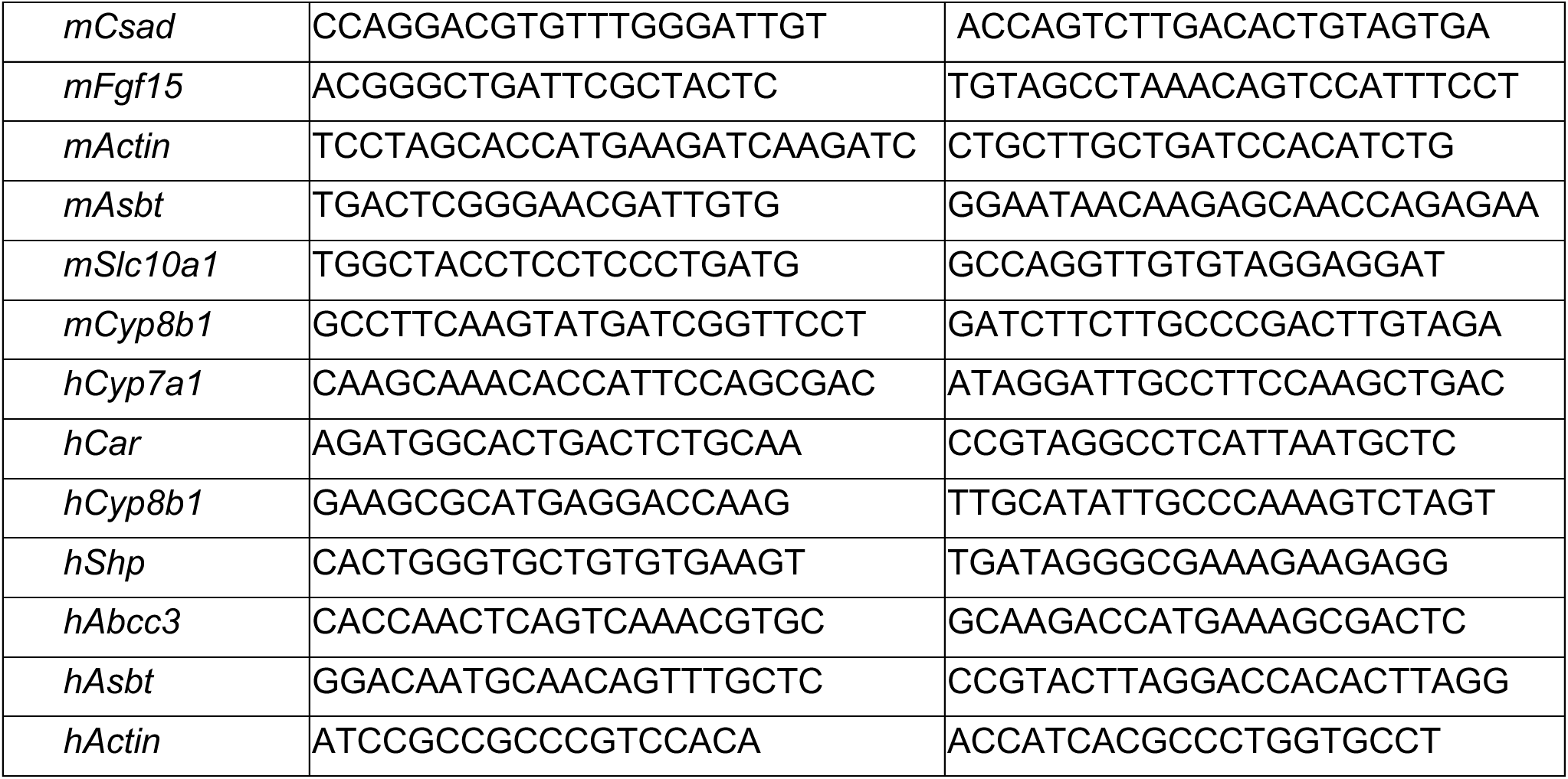

