## Supplemental Figures 1-7 for "AMPK activation prevents hepatocellular carcinoma development through inhibition of HNF4α activity"

### Supplemental Figure 1

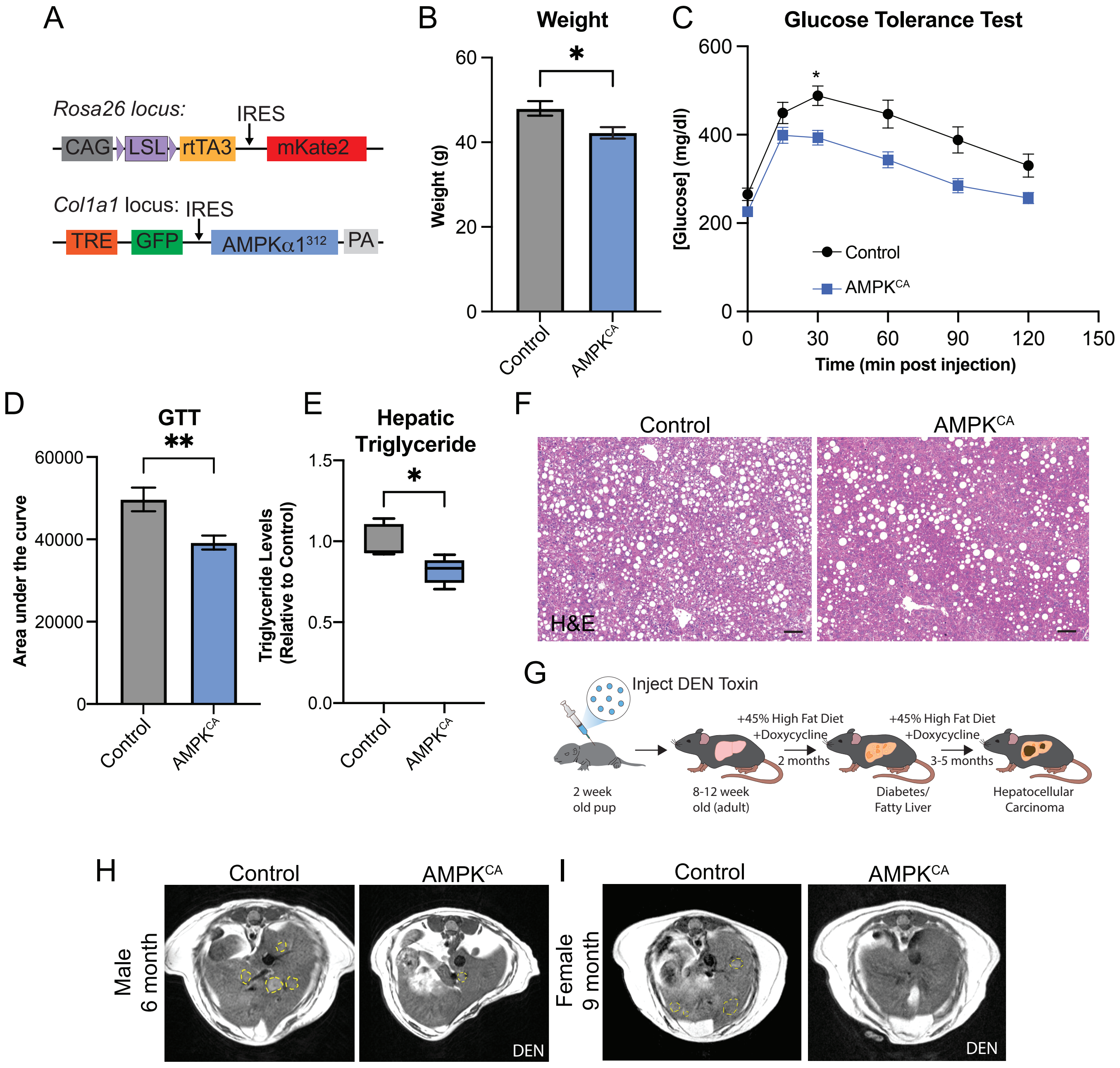


Supplemental Figure 1

A. Schematic of mouse transgenes for expression of mKATE, GFP, and AMPK$^{CA}$. Adapted from Garcia et al, 2017. B. Weight of male control and AMPK$^{CA}$ mice on HFD with doxycycline for 2 months, Welch t-test, ±SEM, n=11-20. C. Glucose tolerance test (1mg/ml Glucose IP; left) and D. area under the curve (right) of control and AMPK$^{CA}$ male mice on HFD with doxycycline for 2 months. n=8-15 ±SEM Welch t-test. E. Quantification of triglyceride levels in liver from female control and AMPK$^{CA}$ mice on HFD with doxycycline for 2 months. n=5 ±SEM. Welch t-test. F. H&E images of control and AMPK$^{CA}$ livers after 7 months on HFD containing doxycycline, Scale bar is 100μm. G. Schematic of DEN-induced HFD HCC model. H. MRI image of male control and AMPK$^{CA}$ livers 6 months post-DEN treatment. I. MRI image of female control and AMPK$^{CA}$ livers 9 months post-DEN treatment *<0.05, **<0.01

Supplemental Figure 2

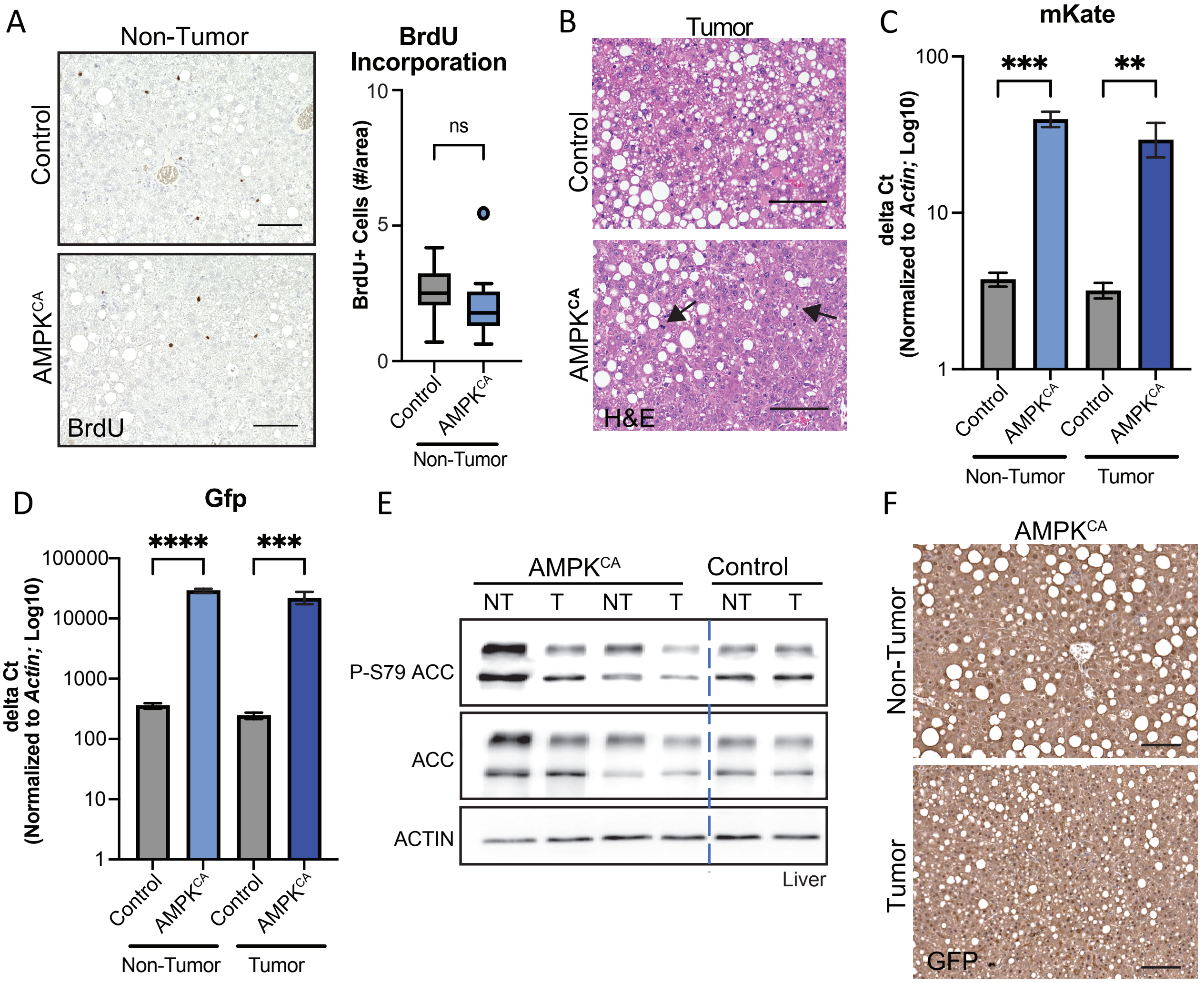


Supplemental Figure 2

A. Immunohistochemical analysis of BrdU incorporation in non-tumor tissue from control and AMPK$^{CA}$ livers following 4 hour BrdU pulse (10mg/kg). Right: Quantification of number of positive cells per area. n=8-11 100μm scale, Welch t-test. B. H&E image of tumors from control and AMPK$^{CA}$ mice matching Fig 2B. Arrows denote mitotic cells. 100μm scale. C. Gene Expression of mKate normalized to *Actin* in non-tumor and tumor tissue. n=5-6, ±SEM, Fisher LSD test. D. Gene Expression of Gfp normalized to *Actin* in non-tumor and tumor tissue. n=5-6, ±SEM, Fisher LSD test. E. Western blot analysis of non-tumor (NT) and tumor (T) tissue. ACTIN as loading control. **<0.01, ***<0.001, ****<0.0001. F. Immunohistochemical analysis of GFP in non-tumor or tumor tissue from AMPK$^{CA}$ livers

### Supplemental Figure 3

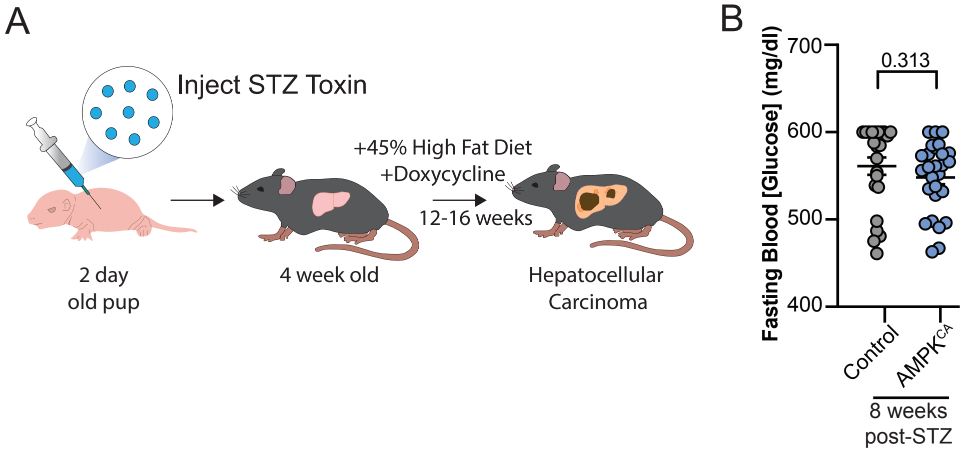


Supplemental Figure 3

A. Schematic of STZ-induced HFD HCC model. B. Fasting blood glucose levels 8 weeks post-STZ treatment. n=23-27. Welch t-test

### Supplemental Figure 4

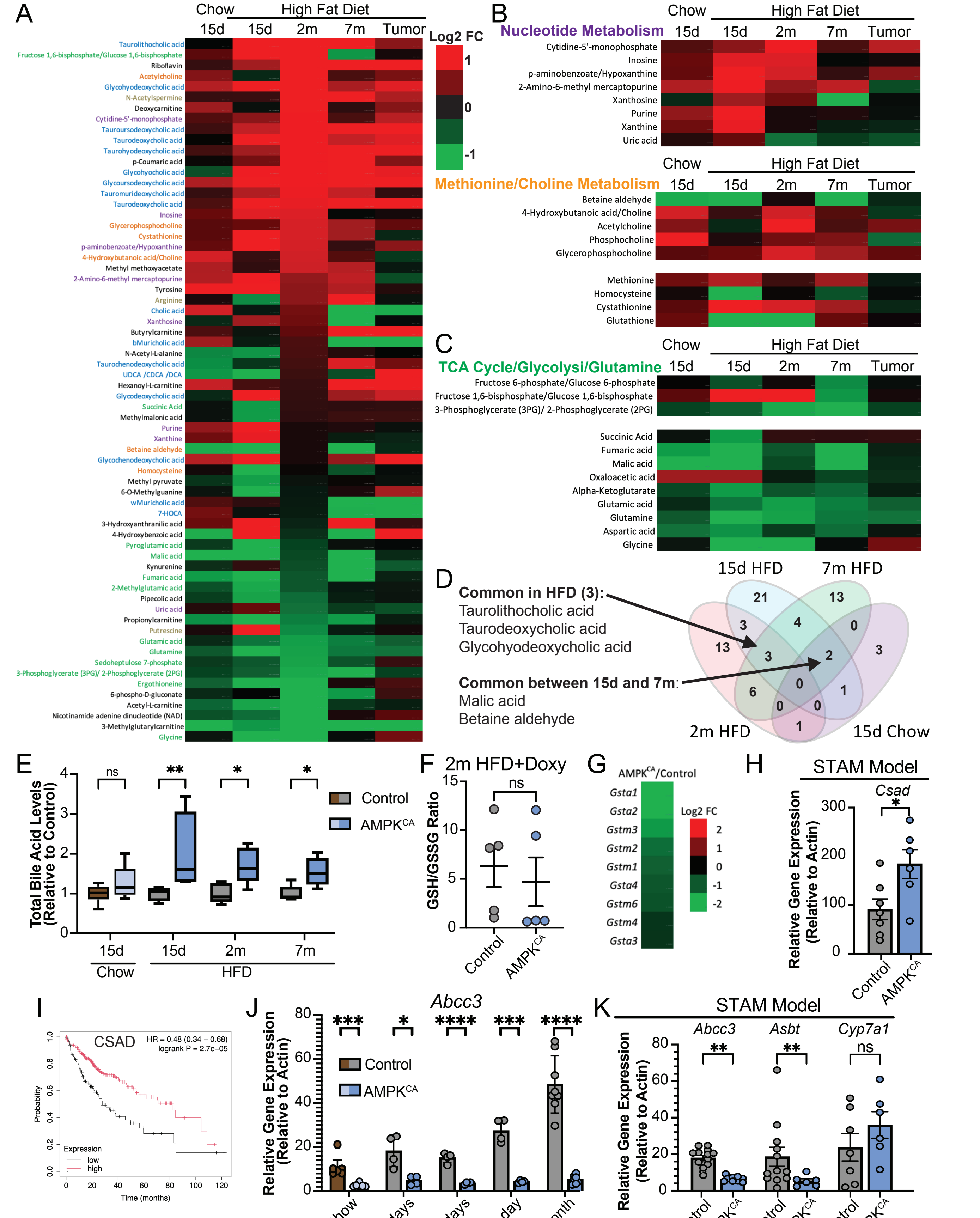


Supplemental Figure 4

A. Heatmap of log2 fold change of metabolites from AMPK$^{CA}$ livers relative to control livers following varying durations of doxycycline treatment (15d, 2m) or in matched non-tumor (7M) and tumors, n=4-6/genotype per timepoint (FC>1.5, p-value<0.05) B. Heatmap of average relative fold change of nucleotide and methionine/choline metabolites from AMPK$^{CA}$ livers relative to control livers following varying durations of doxycycline treatment (15d, 2m) or in matched non-tumor and tumors (7m vs Tumor) n=4-6/genotype per timepoint. C. Heatmap of average relative fold change of TCA cycyle and glutamine metabolites from AMPK$^{CA}$ livers relative to control livers following varying times of doxycycline treatment (15d, 2m) or in matched non-tumor and tumors (7m vs Tumor), n=4-6/genotype per timepoint D. Venn diagram of metabolites changed in AMPK$^{CA}$ livers relative to control livers following varying durations of doxycycline treatment (15d, 2m, 7m) E. Total bile acid levels relative to control following varying durations of doxycycline treatment, 2 month (2m), 7 month (7m) n=4-6. Fisher LSD test, F. Glutathione (GSH) to Glutathione disulfide ratio (GSSG) ratio in livers from mice following 2 month treatment with HFD containing doxycycline, n=5 Welch t-test G. Heatmap of gene expression of Glutatione S Transferase enzymes following 2 month treatment with HFD containing doxycycline H. Quantification of gene expression in control or AMPK$^{CA}$ livers of mice on HFD containing doxycycline following STZ treatment for 20 weeks. n=6-7 Fisher LSD test. I. Survival curve of patients with HCC stratified by high or low expression of CSAD. Kmplot.com. J. Quantification of gene expression in control or AMPK$^{CA}$ livers of mice on chow or HFD containing doxycycline for 3 days, 15 days, or in tumors. n=4-6. Fisher LSD test. K. Quantification of gene expression in control or AMPK$^{CA}$ livers of mice on HFD containing doxycycline following STZ treatment for 20 weeks, n=6-13. Fisher LSD test. ±SEM, *<0.05, **<0.01, ***<0.001, ****<0.0001

### Supplemental Figure 5

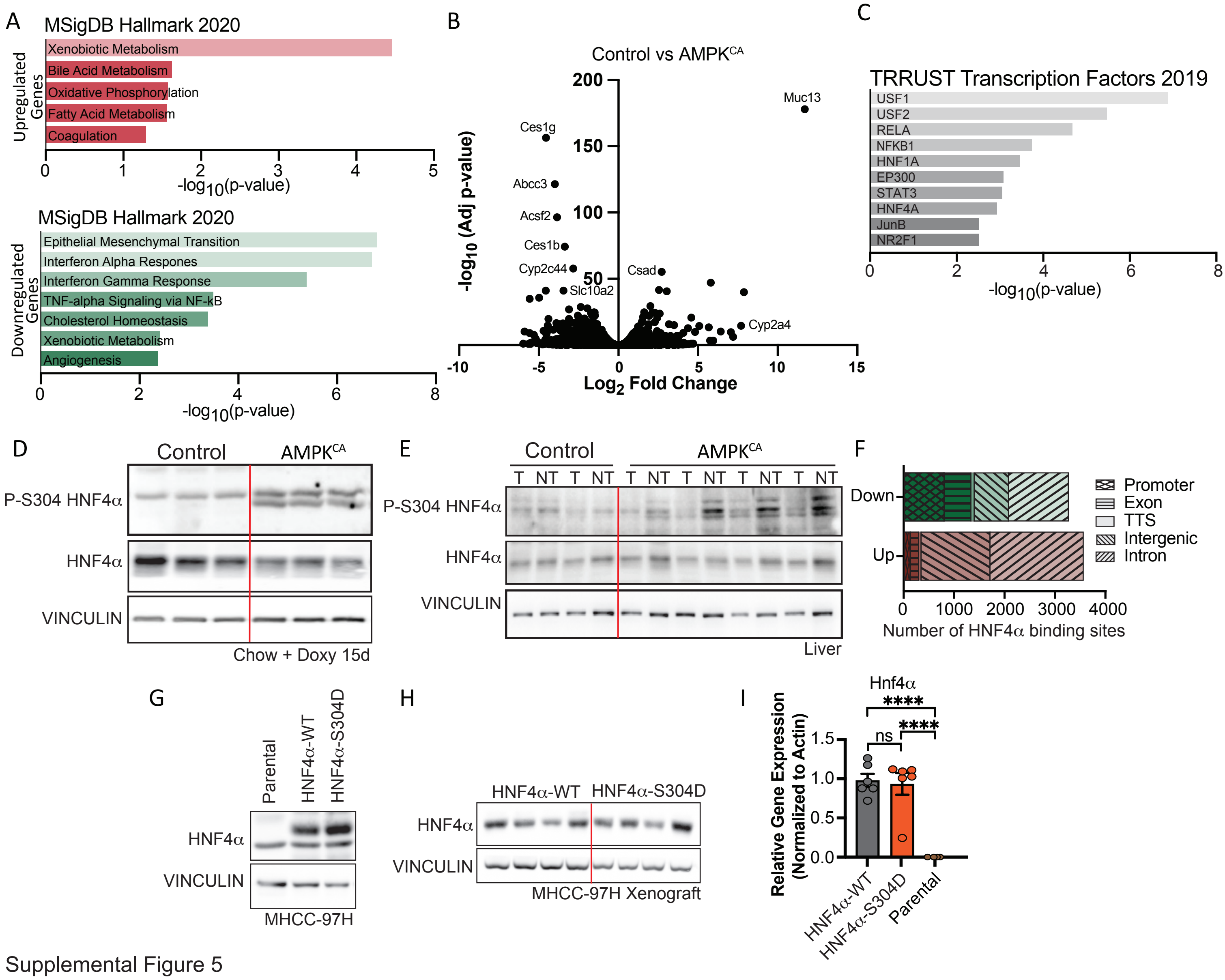


Supplemental Figure 5

A. Hallmarks enriched in upregulated (top) and downregulated (bottom) gene lists from AMPK$^{CA}$ livers compared to control livers, including wild-type mice treated with and without doxycycline or albumin-cre expressing mice in the absence of doxycycline, with a 1.3 fold change with an adjusted p-value less than 0.05. B. Volcano plot of genes changed in AMPK$^{CA}$ livers compared to control livers treated with HFD with Doxycycline with bile acid related genes labeled. C. Transcription factors enriched in AMPK$^{CA}$ mice relative to control mice using previously published gene expression following 2 months of HFD containing doxycycline using Enrichr. D. Western blot of livers following 15d of Chow with doxycycline diet. E. Western blot of tumor (T) and non-tumor (NT) tisse from AMPK$^{CA}$ livers compared to control livers. F. Genomic location and directionality of change in HNF4α binding sites in livers from control and AMPK$^{CA}$ mice following 15 days of HFD containing doxycycline. TTS - Transcriptional termination site. G. Western blot analysis of MHCC-97H cells expressing HNF4α constructs. H. Western blot analysis of xenografts derived from MHCC-97H cells expressing HNF4α construct. I. Gene expression analysis for HNF4α from xenografts derived from MHCC-97H cells expressing HNF4α constructs relative to control MHCC-97H parental cells. n=6 ±SEM ****<0.0001 Fisher LSD test

Supplemental Figure 6

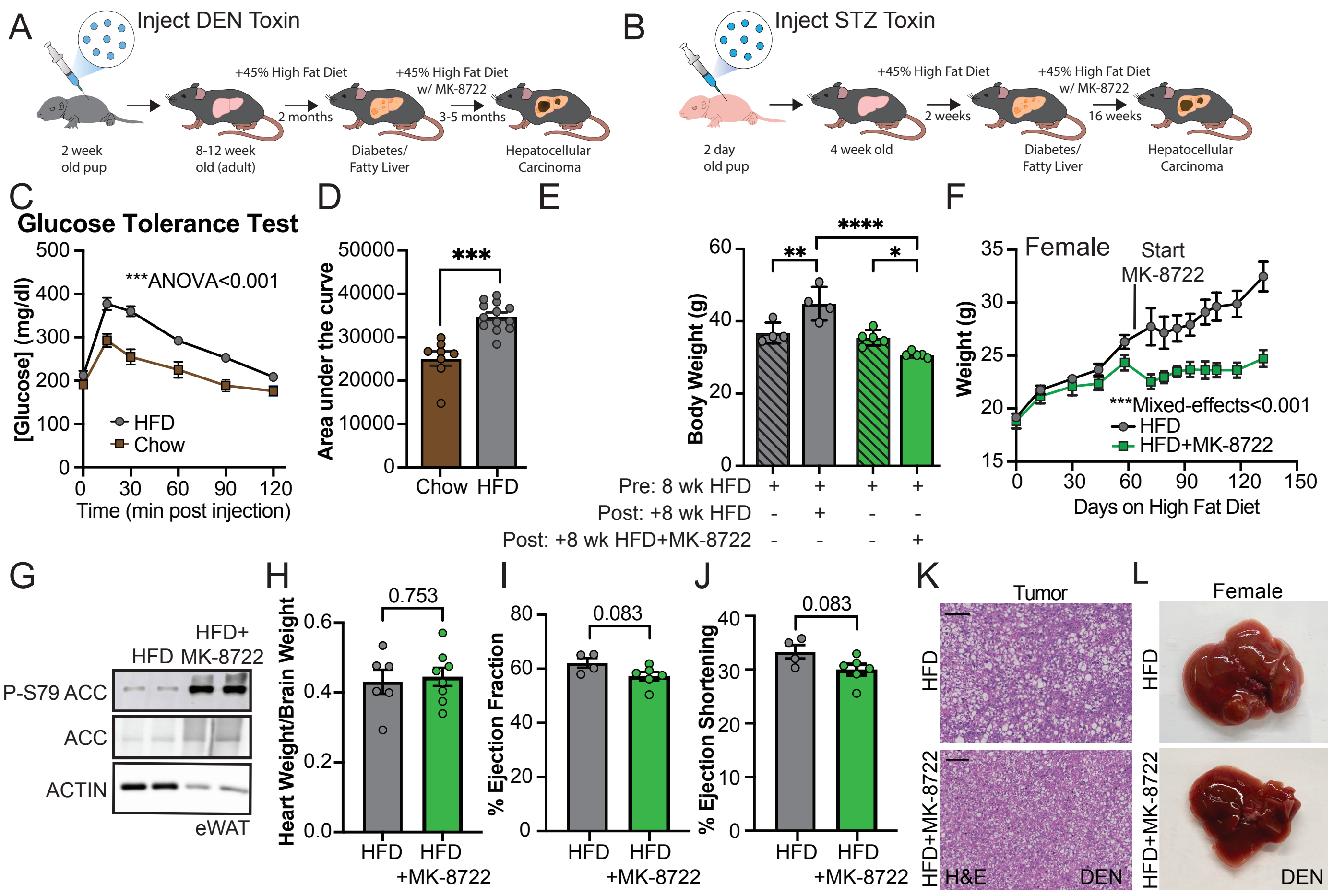


Supplemental Figure 6

A. Schematic of DEN-induced tumor model with MK-8722 treatment B. Schematic of Streptozocin-induced tumor model with MK-8722 treatment C. Glucose tolerance test (1mg/ml Glucose IP; left) of male mice on HFD versus Chow diet for 8 weeks. n=8-13. ±SEM Two-way ANOVA for diet. D. Area under the curve of GTT from male mice fed HFD or Chow diet for 8 weeks. n=8-13 ±SEM Welch t-test. E. Body weight (grams) of male mice fed high fat diet (HFD) for 8 weeks (striped) followed by HFD or HFD+MK-8722 (solid) for 8 weeks, n=4-5 ±SEM Two-way ANOVA. F. Weight Curve of female mice fed HFD for 8 weeks followed by HFD or HFD + MK-8722, n=19 ±SEM Mixed effects analysis. G. Western blot analysis of male mouse epidemal white adipose tissue (eWAT) fed high fat diet (HFD) containing MK-8722 to activate AMPK for 7 days compared to control HFD eWAT. H. Heart weight relative to brain weight of male mice on HFD or HFD+MK-8722 for 5 months. n=6-8 ±SEM Welch t-test. I. Ejection fraction from echocardiogram from male mice injected with DEN and on HFD for 2 month followed by HFD or HFD+MK-8722 for 4.5 months, n=4-6, ±SEM , Welch t-test. J. Ejection shortening from echocardiogram from male mice injected with DEN and on HFD for 2 month followed by HFD or HFD+MK-8722 for 4.5 months, n=4-6, ±SEM Welch t-test. K. Representative H&E image of tumors from DEN-treated male mice on HFD or HFD+MK-8722. Scale bar 100µm. L. Representative whole mount image of female livers fed HFD or HFD+MK-8722 9-months post-DEN treatment. *<0.05, **<0.01, ***<0.001, ****<0.0001

### Supplemental Figure 7

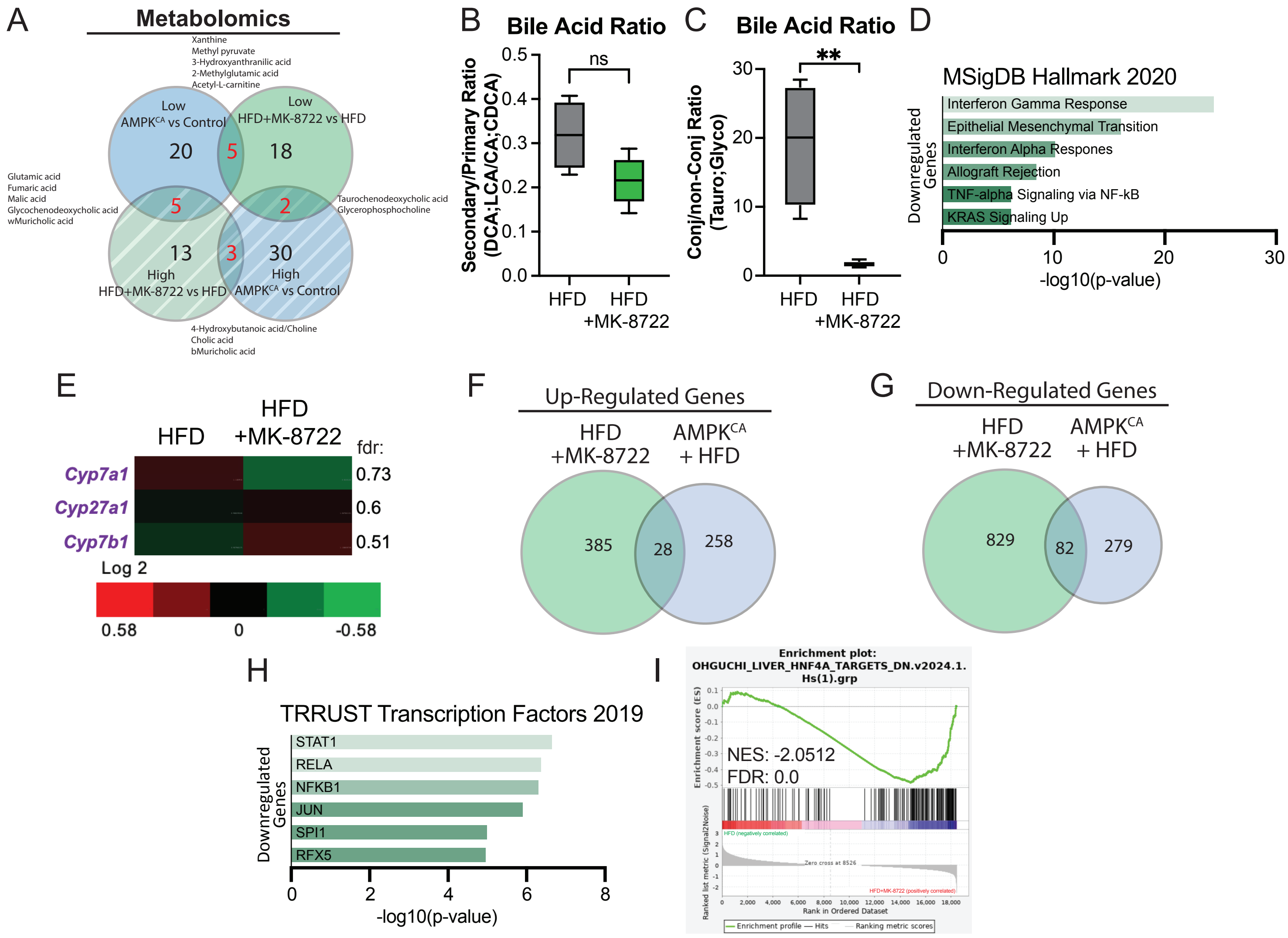


Supplemental Figure 7

A. Venn Diagram of overlapping metabolites changed (low: Solid; high: Striped) between AMPK$^{CA}$ livers and MK-8722 livers relative to control at 2 month. B. Quantification of ratio of secondary to primary bile acids in livers of mice fed HFD or HFD+MK-8722 for 2 months n=4-5, Welch t-test. C. Quantification of ratio of conjugated to non-conjugated bile acids in livers of mice fed HFD or HFD+MK-8722 for 2 months n=4-5, **<0.01 Welch t-test. D. Hallmarks enriched in downregulated gene lists from mice fed HFD+MK-8722 for 2 months relative to control mice fed HFD using Enrichr. E. Heatmap of log2 fold change of genes involved in bile acid synthesis in livers of mice fed HFD+MK-8722 relative to HFD for 2 months with fdr indicated. F. Overlap of genes upregulated upon AMPK$^{CA}$ expression or treatment with MK-8722 in the presence of HFD for 2 months. G. Overlap of genes downregulated upon AMPK$^{CA}$ expression or treatment with MK-8722 in the presence of HFD for 2 months. H. Transcription factors enriched in downregulated genes from mice fed HFD+MK-8722 for 2 months relative to control mice fed HFD using Enrichr. I. Gene set enrichment analysis of genes in mice fed HFD+MK-8722 for 2 months relative to control mice fed HFD
